# A developmental program of early residency promotes the differentiation of divergent uterine NK cell subsets in humans

**DOI:** 10.1101/2025.07.05.663267

**Authors:** Morgan Greene, Rebecca Asiimwe, Emma Wright, Markayla Bell, Daniel Epstein, Brittney Knott, Samantha Fry, Stephanie Clevenger, Holly E. Richter, Aharon G. Freud, Shawn C. Little, Paige M. Porrett

## Abstract

Human uterine natural killer cells (uNKs) are a tissue-resident, innate lymphocyte population that have critical roles in supporting pregnancy health. uNKs derive from circulatory cells in the peripheral blood which immigrate into the endometrium and become resident as they reconstitute the uterine lining after menses. How tissue-resident uterine NK cells arise from blood-based precursor cells is unknown. Here, we identify early tissue immigrants, developmental intermediates, and mature effector states in human endometrium. We also uncover a transcriptional program of TGF-β responsive genes that is upregulated in recent tissue immigrants prior to expression of effector molecules. Differences in TGF-β responsiveness of uNK precursors promote differential expression of divergent effector uNK subsets, resulting in either repression or preservation of cytotoxic effector potential. Collectively, these data suggest a molecular mechanism of tissue-resident uNK maturation that links tissue residency with the acquisition of divergent effector functions in human endometrium.

## INTRODUCTION

Uterine natural killer cells (uNKs) are a tissue-resident, innate lymphocyte population that have critical roles in supporting pregnancy health. The importance of uNKs in pregnancy homeostasis has been demonstrated in both mice and humans, as these cells provide a variety of cytokines which influence spiral artery remodeling^1–5^, trophoblast differentiation^6^, and fetal growth^7^. Disruptions of uNK abundance or function have been associated with a spectrum of reproductive phenotypes and pregnancy complications. In mice, blockade of signals that support uNK development or function results in pregnancy loss, fetal growth restriction, and incomplete transformation of the uterine spiral arteries^8–13^. In humans, alterations of uterine NK abundance or frequency have been associated with pre-eclampsia^14^ and recurrent pregnancy loss^15^. Moreover, pregnant women with certain killer immunoglobulin receptor (KIR) genotypes have an increased risk of pre-eclampsia when the fetus expresses paternal HLA-C alleles of the HLA-C2 group^16^, showcasing the physiologic relevance of uNK interactions with placental trophoblast. Notably, the detrimental effect of this combination of uNK receptors and HLA-C ligands on spiral artery transformation and fetal growth restriction has been shown in humanized mice^12^. Taken together, such studies illustrate the variety of ways that uterine NKs contribute to both physiologic and pathologic pregnancy.

Recent studies of human decidua suggest that the uNK compartment is composed of three subsets of tissue-resident NK cells (i.e., dNK1, dNK2, dNK3 cells) with distinct transcriptional signatures^17^ and surface phenotypes^18^. Comparable subsets have been described in the non-pregnant endometrium by multiple investigators^14,19,20^, suggesting that dNK cells expand during first trimester pregnancy from newly recruited cells from the peripheral blood and/or proliferation of pre-existing tissue-resident uNKs in the endometrium. Importantly, studies of endometrial biopsies and the menstrual blood of uterus transplant recipients suggest that uNKs originate in the periphery at some point in their life cycle, as uNKs express the genotype of the recipient and not the donor^14,21^. These data thus suggest that uNKs derive from circulatory cells in the peripheral blood which immigrate into the endometrium and become resident as they reconstitute the uterine lining after menses.

Although significant progress has been made in our understanding of the molecular mechanisms responsible for the development of tissue residency in human tissues^22–27^, how three subsets of tissue-resident uterine NK cells arise from blood-based precursor cells is unknown. While prior work has suggested that CD39+KIR+ dNK subsets may differentiate from CD39-KIR-dNK cells^28^, no studies to date have included analysis of blood-based precursors to help define the early events or the molecular programs governing uNK differentiation. And while recent studies in mice have identified a key role for TGF-β in the development of uterine tissue-resident NK cells^29,30^, murine dNK cells do not phenocopy human dNK subsets^8,29–32^ and therefore may not provide relevant insights into the differentiation of heterogeneous uNK subsets which exist in humans. Finally, mice have an estrous cycle in lieu of a menstrual cycle^33^ and thus have an altered tissue microenvironment compared to humans. This difference between the reproductive biology of the two species has considerable implications, because tissue regeneration and decidualization are dictated by cyclical changes in ovarian hormones in humans which impact the production of cytokines known to influence uNK survival and maturation^28,34^. There thus exists a significant need to study uterine NK cell differentiation in human tissue samples that are collected from individuals at consistent time points in the menstrual cycle to minimize variation due to differences in the endometrial microenvironment.

In this work, we aimed to construct a comprehensive model of human uterine NK development using multimodal single-cell technologies and analysis approaches that could identify early tissue immigrants, developmental intermediates, and mature effector states in human endometrium. We leveraged new insights into the molecular mechanisms governing early tissue residency in mucosal tissues to identify a transcriptional program of TGF-β responsive genes that is upregulated in recent tissue immigrants prior to expression of the adhesome. This early residency program (ERP) consists of a network of AP-1 and nuclear transcription factors that increase throughout uNK maturation and associate with the expression of TGF-β-induced immunoregulatory cytokine genes in mature decidual-like NK subsets. Moreover, differences in TGF-β responsiveness in the pool of uNK developmental intermediates promote differential expression of the ERP in divergent effector uNK subsets, resulting in either repression or preservation of cytotoxic effector potential. Collectively, these data suggest a molecular mechanism of tissue-resident uNK maturation that links tissue residency with the acquisition of divergent effector functions in human endometrium.

## RESULTS

### Identification of candidate decidual NK precursor cells in human endometrium

To build a developmental framework to describe human uNK differentiation, we first aimed to catalog all potential phenotypic states in human endometrium. We approached this task from the premise that the endometrium itself represents a transitional tissue state that is differentiating into the decidua across the secretory phase^35,36^. We thus expected that endometrial NK cells could exist in a variety of developmental states ranging from recent immigrants from the blood to mature dNK-like cells expressing phenotypes like the decidua. Although there is evidence that endometrial dNK-like cells undergo further differentiation as the endometrium transforms into the decidua after embryo implantation^37,38^, we presumed for the purpose of this work that dNK-like endometrial NK cells would represent the endpoint of our developmental model. To test the hypothesis that immediate precursors of dNK cells sharing dNK phenotypes would be present in human endometrium, we analyzed secretory phase endometrial biopsies (n=8) (Table S1) using flow cytometry. We used a panel of fluor-conjugated antibodies that would identify benchmark decidual phenotypes (i.e., dNK1, dNK2, dNK3 cells)^17^ and label antigens that have been associated with endometrial NK maturation (i.e., KIRs, CD39)^28^ as well as lymphocyte tissue residency across diverse tissues in humans and in mice (i.e., CD49a, CD103)^17,18,20,22,24,26,29,31,39–52^. To this end, we utilized a manual gating strategy which would capture tissue-resident dNK1-like cells (CD49a+CD16-CD103-CD39+KIR+), dNK2-like cells (CD49a+CD16-CD103-CD39-KIR-), dNK3-like cells (CD49a+CD16-CD103+CD39+KIR-), and all possible combinations of these five antigens (Figures 1A and S1A). Secretory phase biopsies were analyzed because eNKs are most abundant at this time point in the menstrual cycle^37,38,53^ and the endometrial microenvironment most closely approximates human decidua. Because this panel of five antigens had the potential to identify 32 phenotypic states (i.e., 2^5^; Table S2), we maximized our ability to capture less abundant phenotypes by concatenating samples for analysis. Overall, we detected 31 of 32 potential phenotypes by manual gating, with similar frequencies of individual phenotypes in all 8 samples (Figures S1B and S1C; Table S2). 57% of endometrial NK (eNK) cells possessed surface phenotypes corresponding to the previously described major dNK subsets (i.e., dNK1, dNK2, and dNK3) (Figures 1A-1B and S1B; Table S2). An additional 28% of eNK cells had surface phenotypes corresponding to other reported but less abundant dNK cells (i.e., alternative trNK), including KIR+ CD103+ trNK3 and KIR+ CD103-CD39-trNK2 (Figures 1A-1B and S1B-S1C; Table S2). Notably, 15% of eNK cells had phenotypes that were not previously described in human decidua^17,18^. This included cells expressing CD49a+CD16+ phenotypes (i.e., double positive cells; “DP”), CD49a-CD16-phenotypes (i.e., double negative cells; “DN”), and CD49a-CD16+ conventional NK cells (“cNK”). Although other investigators have attributed these phenotypes to blood contamination of decidual preparations^17,18^, we opted to include these phenotypes in further analyses as these cells may represent recent endometrial immigrants (i.e., founder NK cells that have not yet upregulated CD49a) or other developmental intermediates. Altogether, these results suggested that while ∼60% of eNKs express major decidual-like phenotypes as suggested in prior work^17,18,28^, almost half of NK cells in human endometrium have been less well characterized and may represent a range of dNK precursors or alternative phenotypes.

**Figure 1.**
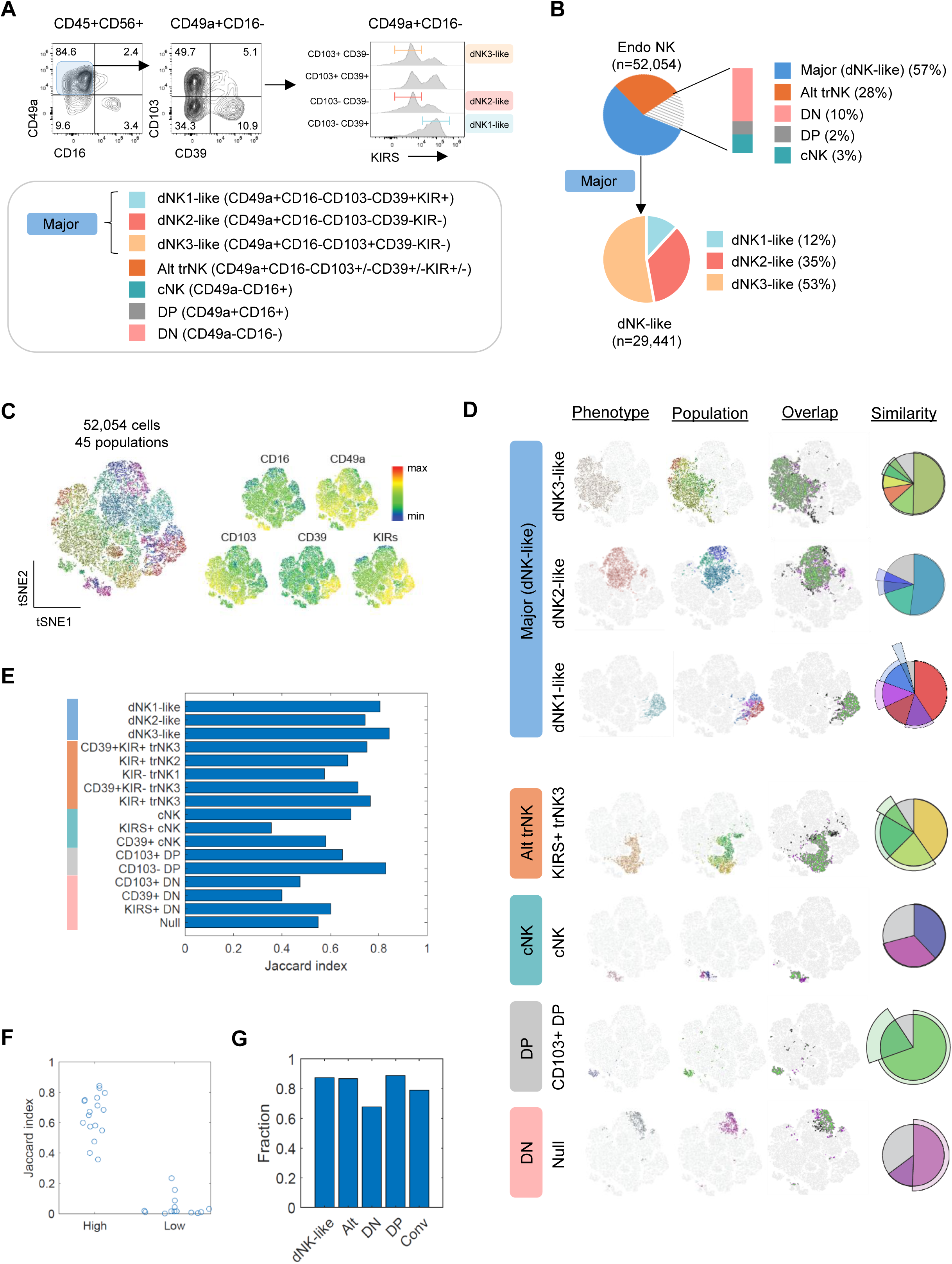
NK phenotypic states in secretory phase human endometrium. (A-G) Human endometrial biopsies (n=8) were digested and analyzed with flow cytometry. Live, CD45+CD3-CD14-CD56+ singlets were gated in each sample and then concatenated for downstream analysis. (A-B) Phenotypes (A) and frequency (B) of major phenotypic groups of CD56+ endometrial NKs by manual gating. See Table S2 for descriptions of all 32 phenotypes and Figure S1 for enumeration of high confidence phenotypes. (C-G) FlowSOM analysis of concatenated data. (C) tSNE representation of 45 FlowSOM-generated populations (left) and expression of individual surface proteins (right). (D) Comparison of manually gated phenotypes and FlowSOM-generated populations. Seven of 17 high confidence phenotypes are shown. See Figure S1 for other high confidence phenotypes. Similarity polar area charts quantify overlap of manually gated phenotypes (circle) and FlowSOM populations (colored wedges). Populations that contribute to the phenotype are those which have a majority of cells belonging to the phenotype. Area outside of the circle (“pie crust”) represents frequency of a population that belongs to a different phenotype. Gray wedge inside the pie represents the percent of cells of the designated phenotype that belong to populations where they represent the minority of cells. (E) Quantification of overlap (Jaccard Index) for 17 high confidence phenotypes. (F) Distribution of the Jaccard Index for high confidence (n=17) and low confidence (n=14) phenotypes. (G) Fraction of high confidence cells within each major phenotypic group. High confidence cells are defined as cells assigned to the same phenotype by both manual gating and FlowSOM (i.e., inside the pie, (D)). Low confidence cells are cells that are assigned to one phenotype by manual gating and one phenotype by FlowSOM (i.e., pie crust or gray pie wedge (D)).

We next considered whether assumptions around the existence of certain phenotypic states and population hierarchies might have biased our phenotypic characterization of endometrial NK cells. Examples of such assumptions include that all tissue-resident populations would express CD49a or that surface proteins associated with maturity in peripheral blood NK cells (i.e., KIRs) would indicate the same developmental state in tissue-resident NK cells. To combat bias in our analytic approach and improve our confidence in the existence of less abundant phenotypes, we analyzed our data using an unsupervised methodology (i.e., FlowSOM^54^) with the expectation that such an approach would validate our gating strategy if FlowSOM identified similar patterns of antigen co-expression. To assure capture of nuanced phenotypes, we intentionally over-clustered the concatenated data and found 45 populations given variable expression of all five antigens (Figure 1C). We reasoned that overclustering to create FlowSOM populations would have at least three effects: (1) large clusters representing a single phenotype would be broken into multiple FlowSOM-generated populations, but as long as such clusters were reasonably well-separated from the remainder of cells, all such populations would consist of a majority of that phenotype; (2) small clusters with few cells representing a single phenotype would represent the majority of at least one FlowSOM population if that population were well-separated from the remainder of cells; and (3) unreliable or low-confidence phenotypes consisting of few cells would be distributed among larger populations and never represent the majority of any FlowSOM population.

With this reasoning, we used FlowSOM to classify a phenotype as high confidence based on whether FlowSOM generated population(s) where the majority of cells consisted of that phenotype. Using this criterion, we found 17 high confidence and 15 low confidence phenotypes (Figures 1C-1F and S1D-F). High confidence phenotypes included all three CD49a+CD16-dNK-like phenotypes as well as all five CD49a+CD16-alternative trNK phenotypes (Table S2). High confidence phenotypes also included 3 of 8 CD49a-CD16+ cNK phenotypes, 2 of 8 CD49a+CD16+ double positive phenotypes, and 4 of 8 CD49a-CD16-double negative phenotypes (Table S2). Altogether, these results suggest that about 40% of human endometrial NK cells possess phenotypes that have not yet been well characterized in human decidua.

Although 98.7% of NK cells in the endometrium were assigned to one of the 17 high confidence phenotypes (Figure S1D), we noted that a fraction of cells within a phenotype were found in FlowSOM populations not assigned to that phenotype. Because these cells grouped with different sets of cells in the two analysis methods, the phenotypes of these individual cells are uncertain despite confidence in the phenotype of neighboring cells. These lower confidence cells tended to be found at the interfaces between phenotypes (Figure 1D) and belonged to every phenotypic group, including dNK-like major phenotypes (Figures 1E and 1G). For example, ∼15% of cells assigned to one of the three dNK-like phenotypes were found in FlowSOM populations that were not composed of the majority of those phenotypes (Figure 1G). The disagreement of the two analysis methods likely arises from the continuum of expression values of these cell surface proteins and suggests that the actual phenotypes of these cells do not align well with a simple binary classification scheme. It is therefore reasonable to conclude that these cells of uncertain phenotype represent cells in transitional states between two or more phenotypes. After the addition of these uncertain cells not captured by the benchmark classification scheme^17^, we found the fraction of unaccounted cells increases to 49%. Altogether, we conclude that at least 40% of human endometrial NK cells exist in phenotypic states outside of classically described decidual NKs.

### Single-cell analysis of NK phenotypes and transcriptomes reveals abundance of putative transitional states in human endometrium

We next aimed to identify human uNK differentiation states using CITE-seq analysis, which allowed direct linkage of surface phenotypes with transcriptomic states. We aggregated NK cells from CD45+ immune cells enriched from three additional secretory phase endometrial biopsies (Figure S2A). We first confirmed the reproducibility of phenotype capture across methodologies and samples by comparing the surface phenotypes of samples analyzed by CITE-seq with those analyzed by flow cytometry. We detected all high confidence 17 phenotypes of endometrial NK cells by examination of the relevant surface proteins (Figures 2A and 2B; Table S2). Moreover, the frequency of these phenotypes was similar overall between individuals analyzed by flow cytometry (n=8) vs. CITE-seq (n=3) (Figure S2B), thus confirming that dNK-like cells represented approximately 50% of cells in these samples according to surface phenotype.

**Figure 2.**
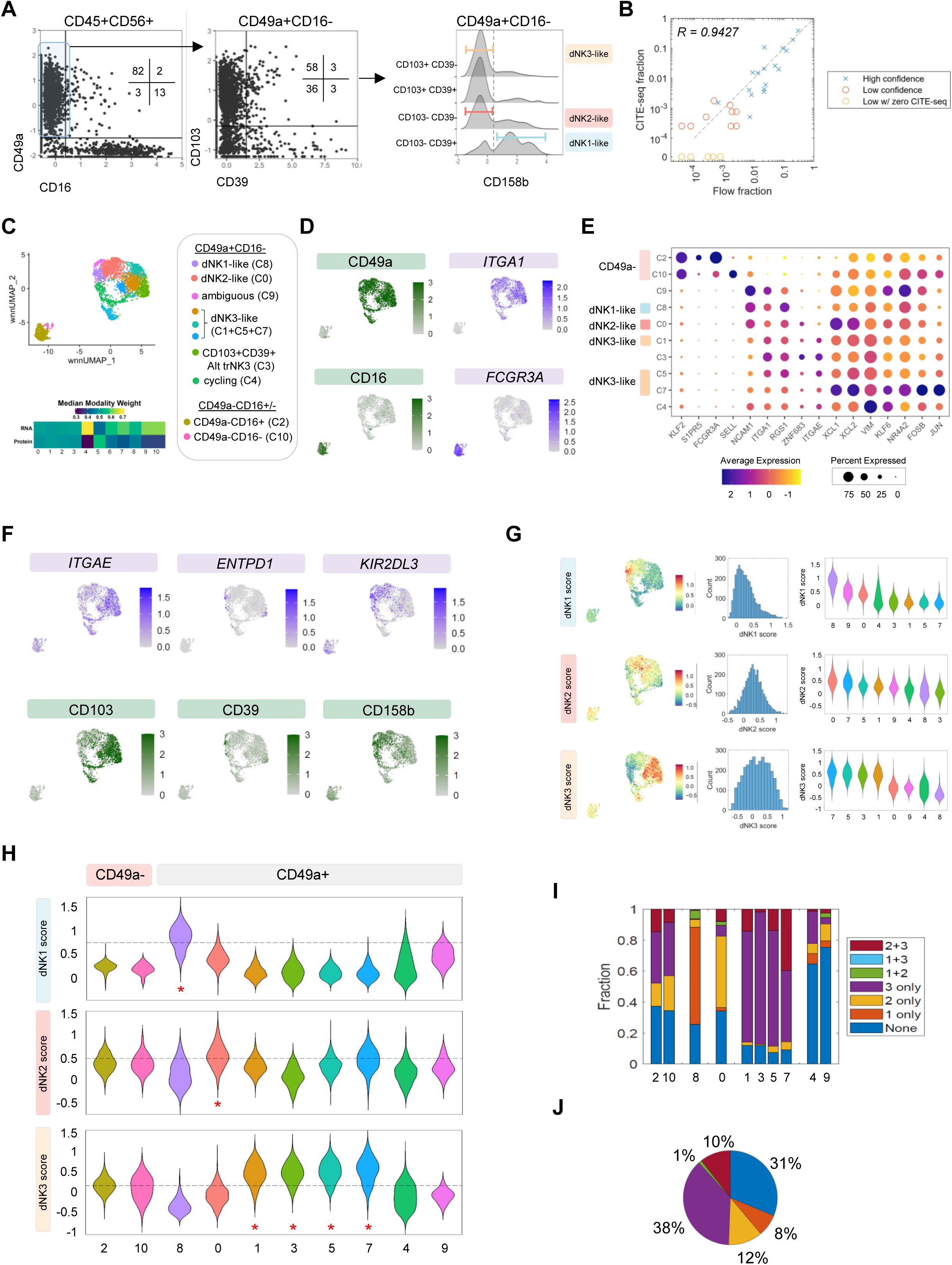
NK transcriptomic states in secretory phase human endometrium. (A-J) CITE-seq analysis of 3,957 CD56+ NK cells selected from 13,748 CD45+ immune cells enriched from secretory phase biopsies (n=3). Data were aggregated for analysis. (A-B) Surface phenotype (A) and quantification (B) of endometrial NK cells analyzed by CITE-seq. Cells were selected using a manual gating approach to compare with flow cytometry results in Figure 1. Comparison of phenotype frequency between the two datasets and approaches (B). (C-E) Weighted nearest neighbor (wnn)UMAP projection of endometrial NK cells reveals two major groups of endometrial NK cells distinguished by reciprocal expression of *ITGA1* (CD49a) and *FCGR3A* (CD16). (F-J) Annotation of endometrial tissue-resident subsets. (F) Expression of marker genes and proteins associated with decidual NK subsets. (G) Continuous expression of decidual NK reference gene signatures illustrates the challenges associated with dNK-like subset identification. (H) Cluster assignment of CD49a+ endometrial trNK. The threshold for each dNK reference signature is shown as a dotted line. Red asterisks indicate cluster(s) meeting criteria for assignment to a given signature. *ITGA1*-cells are shown for comparison. Note four clusters do not meet criteria for assignment. (I-J) Cell-type assignment of *ITGA1*+ endometrial NKs based on dNK signature expression above or below the optimized threshold. Frequency of cells in each cluster (I) or total cells (J) with no assignment (dark blue), single assignment (orange [dNK1-like], yellow [dNK2-like], purple [dNK3-like]), or multiple assignments (light blue, red, green) is shown (I).

Having established that these samples possessed equivalent phenotypic states, we then performed weighted nearest neighbor clustering^55^ of endometrial NK cells to identify uNK cells according to shared phenotypes and transcriptional profiles (Figures 2C and S2C). Variation in both cellular phenotypes and transcriptomes contributed to the cluster assignments for most cells in the samples, although one cluster of cycling NK cells (cluster 4) had a higher RNA modality weight (Figures 2C and S2D). After exclusion of a cluster of NKT cells (Figure S2D), we found that two major groups of endometrial NK cells were distinguished by reciprocal expression of CD49a (*ITGA1*) and CD16 (*FCGR3A*) among other variable genes and proteins (Figures 2D-2F and S2E-G). In line with our flow cytometry results, CD49a+CD16-cells were the most abundant subset of uNKs and were designated as tissue-resident NK cells (trNK) given their expression of integrins (i.e., CD49a, CD103) as well as other genes associated with tissue residency (*RGS1, ZNF683*) (Figures 2E and 2F). Expectedly, subsets of CD49a+CD16-trNKs corresponded to previously described dNK phenotypic states based on expression of CD103 (*ITGAE*) and CD39 (*ENTPD1*)^17^ (Figure 2F). The next most abundant major group were CD56^dim^CD16+ cells (Cluster 2), which were CD49a-negative and expressed genes associated with circulation such as *KLF2* and *S1PR5* (Figures 2E and S2F). Likewise, a minor cluster of CD49a-CD16-double negative cells (C10) which clustered in proximity to the CD49a-CD16+ cells also expressed *KLF2* and *S1PR5*. Taken together, these CD49a-clusters possessed transcriptomic profiles and surface phenotypes which mirrored CD56^dim^CD16+ and CD56^bright^CD16-peripheral blood NK cells. Overall, these results recapitulated the results of our flow cytometry findings and illustrated the utility of CD49a and CD16 in distinguishing major groups of uterine NK cells, including tissue-resident CD49a+ trNK cells.

We next focused our analysis on identifying the trNK clusters and cells which might correspond to mature dNK-like cells. To this end, we utilized reference gene signatures for dNK1, dNK2, and dNK3 cells derived previously^17^ and calculated module scores for all three signatures for each cell in the dataset (Figure 2G). Because of broad and overlapping distributions of the module scores within a given signature, we applied a thresholding and odds ratio maximization procedure to assign one or more clusters to dNK signatures (see Methods). In brief, for each signature, we first ranked clusters in descending order of their mean score, generating an ordered list of candidate clusters that might be assigned to a signature. We assumed that at least one cluster must correspond to the cell type represented by each of the three signatures, or in other words, that cells of the type represented by a signature are present in sufficient numbers to form a cluster. Starting from the first cluster and subsequently adding clusters further down the list, for each cluster combination we found a threshold value for every reference dNK signature that jointly maximized (1) the fraction of cells above the threshold and included in candidate cluster(s) assigned to that cell type, and (2) maximized the fraction of cells below the threshold and not in the candidate cluster(s). Thresholding allowed the calculation of an odds ratio quantifying the performance of the cluster-signature assignments: a relatively high odds ratio indicates that most cells in clusters assigned to that cell type are above the threshold. More than one cluster was assigned to a signature if assignment of additional clusters increased the odds ratio (Figure S2H). Based on this process, we assigned cluster 8 to dNK1-like cells, cluster 0 to dNK2-like cells, and clusters 1, 3, 5, and 7 to dNK3-like cells (Figure 2H). Collectively, approximately 26% of cells were not assigned to a decidual NK-like state, including both clusters of CD49a-cells (14%) and two clusters of CD49a+ cells (12%).

The assignment of cells based on clustering, however, ignores dispersion and overlap of signature scores between the clusters, which may lead to misassignments and fails to identify potential shared states. For example, approximately 15% of cells in cluster 9 were above the threshold for the dNK1 signature but were not assigned as dNK1 cells. Conversely, 30% of cluster 8 cells were below the threshold for the dNK1 signature but were assigned as dNK1 cells using this approach. We therefore examined the fraction of cells in each cluster above one or more thresholds to quantify the heterogeneity of cell states within a given cluster (Figure 2H). This approach identified that almost 1/3 of endometrial NK cells did not meet criteria to be classified as dNK-like cells, while 11% of cells were above two thresholds (Figures 2I and 2J). Altogether, these results suggest that about 40% of cells could not be assigned to one and only one dNK-like state. We speculate that such cells could represent transitional states not previously described in decidua (i.e., unassigned) or transitional states between dNK-like cells (e.g., cells that have >1 assignment).

### Founder NK cells and developmental intermediates revealed through comparison of peripheral blood and human endometrium

As our phenotypic and transcriptional analyses collectively indicated that many human endometrial NK cells could not be assigned to dNK-like states, we next aimed to identify unassigned or ambiguous cells and understand their developmental relationships with dNK-like cells. To identify cell states along the early spectrum of tissue-resident NK differentiation, we hypothesized that cells with subtle shifts of the transcriptome would emerge in an aggregated transcriptional analysis of peripheral blood NKs (pbNK) and endometrial NK cells (eNK). To this end, we analyzed CD45+ cells sorted from an endometrial biopsy and a matched peripheral blood sample (Figure S3A) from one individual to minimize noise due to menstrual cycle variation and/or individual genetic differences. As we did not capture many hematopoietic stem cells or innate lymphocyte cell precursors (ILCPs) in this sample (Figure S3A-C), we opted to interpret these data using a parsimonious model of trNK development where pbNKs entered the endometrium and thus were the source of tissue-resident eNKs.

Among 2,495 NK cells, we found little transcriptional overlap between pbNKs and eNKs (Figure 3A), and pbNKs did not co-cluster with most endometrial NK cells (Figure 3B). Upregulation of tissue resident genes (i.e., *ITGA1*, *RGS1*, *CD69*) and downregulation of genes associated with circulation in eNKs (i.e., *FCGR3A*, *KLF2*) contributed to the segregation of pbNK and eNK (Figures 3C-D and S3D). After excluding a group of NKT cells (C6), we identified two clusters of *ITGA1^lo^* NK cells in the peripheral blood which corresponded to CD56^dim^CD16+ cells (i.e., pb_CD56^dim^ (C2)) and CD56^bright^CD16-cells (i.e., pb_CD56^bright^ (C7))^56^ (Figures 3D and 3E). In accord with our prior CITE-seq (Figure 2) and flow cytometry analyses (Figure 1), we found *ITGA1^lo^* endometrial NK cells that co-clustered with pbNKs in clusters 2 and 7 (Figures 3A-B). To our surprise, however, we found that eNKs that co-clustered with peripheral blood CD56^dim^CD16+ (C2) and CD56^bright^CD16-cells (C7) expressed genes associated with tissue residency (Figures 3D and 3F), albeit at a lower level than other *ITGA1*+ endometrial NK clusters (Figure 3D). These results suggested to us that *ITGA1*^lo^ eNKs in clusters 2 and 7 may represent founder NK cells and not blood contaminants. To confirm that these putative founder NK cells were not unique to one individual, we replicated this analysis in matched blood and endometrium from another individual with similar findings (Figure S3E-I). We also revisited our CITE-seq analysis of three endometrial biopsies (Figure 2) and noted that CD49a-CD16+ (C2) and CD49a-CD16- (C10) cells expressed several genes associated with tissue residency (i.e., *NR4A2*, *JUN*, *FOS*) that were upregulated in *ITGA1^lo^* endometrial NK cells versus *ITGA1^lo^* peripheral blood NK cells (Figures 3F and S3D; S3H-I), suggesting that CD49a-negative cells in the endometrium did not represent a homogenous group of circulating cells. In our matched blood and endometrial analyses, we further noted that cluster 2 endometrial NK cells downregulated the CD56^dim^CD16+ conventional NK signature to such an extent that the cluster of C2 endometrial NK cells did not meet the cNK signature threshold (Figure 3E). Meanwhile, C7 eNKs were able to maintain expression of genes associated with CD56^bright^CD16- pbNK cells (i.e., *SELL*, *TCF7*, *IL7R*). Altogether, these data suggest that *ITGA1*-negative endometrial NK cells are a heterogenous population of recent endometrial immigrants which may originate from either CD56^bright^CD16- or CD56^dim^CD16+ pbNK cells.

**Figure 3.**
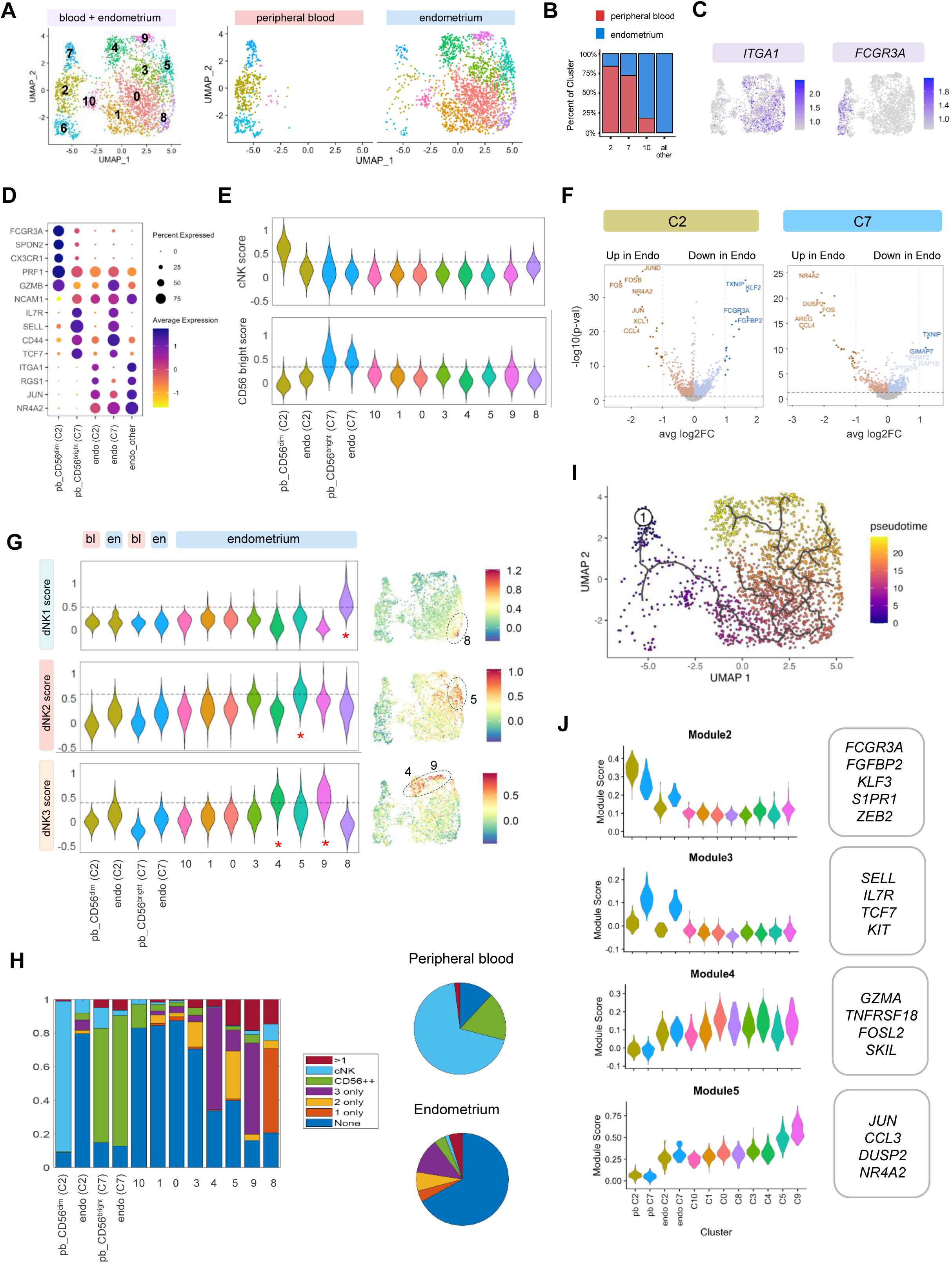
Identification of founder, transitional, and mature dNK-like states among human endometrial NK cells. (A-J) scRNA-seq of 2,495 natural killer cells re-clustered from endometrial and peripheral blood CD45+ immune cells (see Figure S3). Samples were matched from one individual and collected within 24h during the secretory phase of the menstrual cycle. eNK = endometrial NK; pbNK = peripheral blood NK. (A) UMAP embeddings of endometrial and peripheral blood NK cells, shown in aggregate (left) or split by sample (right). (B) Frequency of pbNK and/or eNK cells in each cluster. (C) Reciprocal expression of *FCGR3A* and *ITGA1* demarcates circulating peripheral blood NK cells from tissue-resident endometrial NK cells. (D-F) Identification of founder NK cells in human endometrium. Expression of tissue-resident genes by endometrial NK cells which cluster with peripheral blood CD56^dim^CD16+ (conventional NK; cNK) (C2) and peripheral blood CD56^bright^CD16- (C7) populations (D). Note that C7 eNKs and pbNKs express marker genes associated with CD56^bright^CD16-cells (i.e., *SELL*, *IL7R*) (D&E). In contrast, C2 endometrial NK cells which co-cluster with C2 peripheral blood NK cells have poor expression of cNK markers such as *FCGR3A*, *PRF1*, and *GZMB* and have reduced expression of cNK signature (D&E). Tissue-resident genes which are differentially upregulated in endometrial vs. peripheral blood NK cells are shared between eNK cells which co-cluster with peripheral blood cells (F). (G-H) Identification of transitional and mature dNK-like clusters (G) and cells (H) in human endometrium. Mature dNK1-like, dNK2-like, and dNK3-like clusters are identified based on expression of dNK reference signatures (C8, C5, and C4+C9, respectively, indicated by a red asterisk) (G). However, five endometrial NK clusters (C2, C10, C1, C0, and C3) are unassigned (G) as a minority of cells in these clusters meet thresholds for any of the five signatures (cNK, CD56bright, dNK1, dNK2, dNK3) (H). (I) Trajectory analysis places transitional clusters between founder NK cells (C7) and mature dNK-like clusters. (J) Gene modules that are downregulated and upregulated between founder NK cells (C7) and mature dNK-like clusters (C8 [dNK1-like], C5 [dNK2-like], C4+C9 [dNK3-like]).

We next aimed to identify clusters of dNK-like cells among the *ITGA1*+ endometrial trNKs. Using our thresholding approach for cluster-based assignment, we identified two subsets of trNK3 cells (C4+C9), one subset of dNK2-like cells (C5), and one subset of dNK1-like cells (C8) among 8 clusters of *ITGA1*+ cells (Figure 3G). Half of *ITGA1*+ clusters did not meet criteria to assign as dNK-like (C10, C0, C1, C3), and >70% of cells within each of these clusters could not be assigned to any mature signature (pbCD56^bright^, pbCD56^dim^, or dNK-like) (Figure 3H). In contrast, the frequency of unassigned cells in dNK1-like and dNK3-like clusters ranged from ∼10-30% (Figure 2H), suggesting that unassigned cells might represent transcriptionally undefined transitional states. Nevertheless, we still noted considerable heterogeneity even within mature dNK-like clusters, as many cells were unassigned or met criteria for several signatures (Figure 3H). Among dNK-like clusters (C4, C5, C9, C8), we noted that most cells within the cluster designated as dNK2-like (C5) did not meet the threshold for dNK2-like assignment. In other words, most cells in C5 were unassigned, assigned to dNK3-like cells, or met criteria for multiple states (Figure 3H). This is consistent with prior observations in other endometrial samples (Figures 2I and 2J) and was reflected by the fact that the dNK2 signature had the lowest odds ratio by an order of magnitude (Fig S2H). Taken altogether, these data suggest that only 26% of NK cells in human endometrium represent previously described transcriptional states found in either peripheral blood or decidua.

To test the hypothesis that clusters C10, C0, C1, C3 represented cells transitioning from recent immigrants to mature states, we performed trajectory analysis. Although our prior analysis suggested there might be two populations of founder NK cells, we elected to set cluster 7 as the root of our trajectory given the less mature status of pbCD56^bright^ cells and their stem-like features (i.e., expression of *TCF7*, *SELL*).^57^ In line with our hypothesis that clusters C10, C0, C1, C3 represented transitional cells, the trajectory passed through these clusters to end in clusters designated as dNK-like cells (Figure 3I). Notably, multiple trajectories stemmed from within C0 & C1, suggesting multiple potential differentiation paths for eNKs. Subsequent analysis of the gene modules that changed across the trajectory (Figures 3J and S3J) revealed that genes associated with CD56bright cells (*SELL, TCF7*), CD56dim cells (*FCGR3A, FGFBP2*), and genes associated with circulation (*S1PR1, KLF3*) were reduced in differentiating endometrial *ITGA1*+ trNKs (Figure 3J). Conversely, we found increased expression of genes associated with lymphocyte residency programming in modules 4 and 5 (*FOSL2, NR4A2, JUN*) as pseudotime progressed from putative founder NK cells in C2 and C7 towards mature dNK-like cells in clusters 5 and 9 (Figure 3J). Collectively, these results suggest a developmental model whereby pbNK subsets seed the endometrium to become *ITGA1^lo^* founder NK cells which give rise to developmental intermediates and progressively differentiate into mature dNK-like subsets.

### Conserved transcriptional program of early residency is upregulated from founder to mature human endometrial trNK cells

Having identified populations of putative founder NK cells in human endometrium, we compared these cells with their peripheral blood counterparts to test the hypothesis that genes upregulated in *ITGA1^lo^*eNK cells would represent the earliest transcriptional changes associated with tissue residency. Although our prior analysis suggested that either CD56^bright^CD16- or CD56^dim^CD16+ pbNK cells might give rise to endometrial founder NK cells given the expression of genes associated with tissue residency in either population (Figures 3D and 3F), we theorized that endometrial NK cells co-clustering with CD56^bright^CD16-pbNK cells would represent the primary population contributing to the pool of developmental intermediates. We thus compared C7 eNK cells with C7 pbNK cells and found upregulation of a variety of immediate early response genes associated with nuclear receptor (i.e., *NR4A2*) and AP-1 transcription factor (i.e., *JUN, FOSB*) families in C7 eNKs vs. pbNKs (Figure 4A). As these genes are upregulated at early time points after the entry of CD8 resident memory T cells (T_RM_) in tissues of mice,^45^ we tested the hypothesis that this CD8 T_RM_ signature would be enriched in genes upregulated in cluster 7 eNK cells versus cluster 7 pbNK cells. Consistent with our hypothesis that cluster 7 eNKs represented early immigrant founder NK cells, we found statistically significant enrichment of the CD8 T_RM_ signature in differentially expressed genes upregulated in cluster 7 endoNKs (Figure 4B). We thus designated these 80 CD8 T_RM_ genes enriched in C7 eNK cells as the early residency program (ERP).

**Figure 4.**
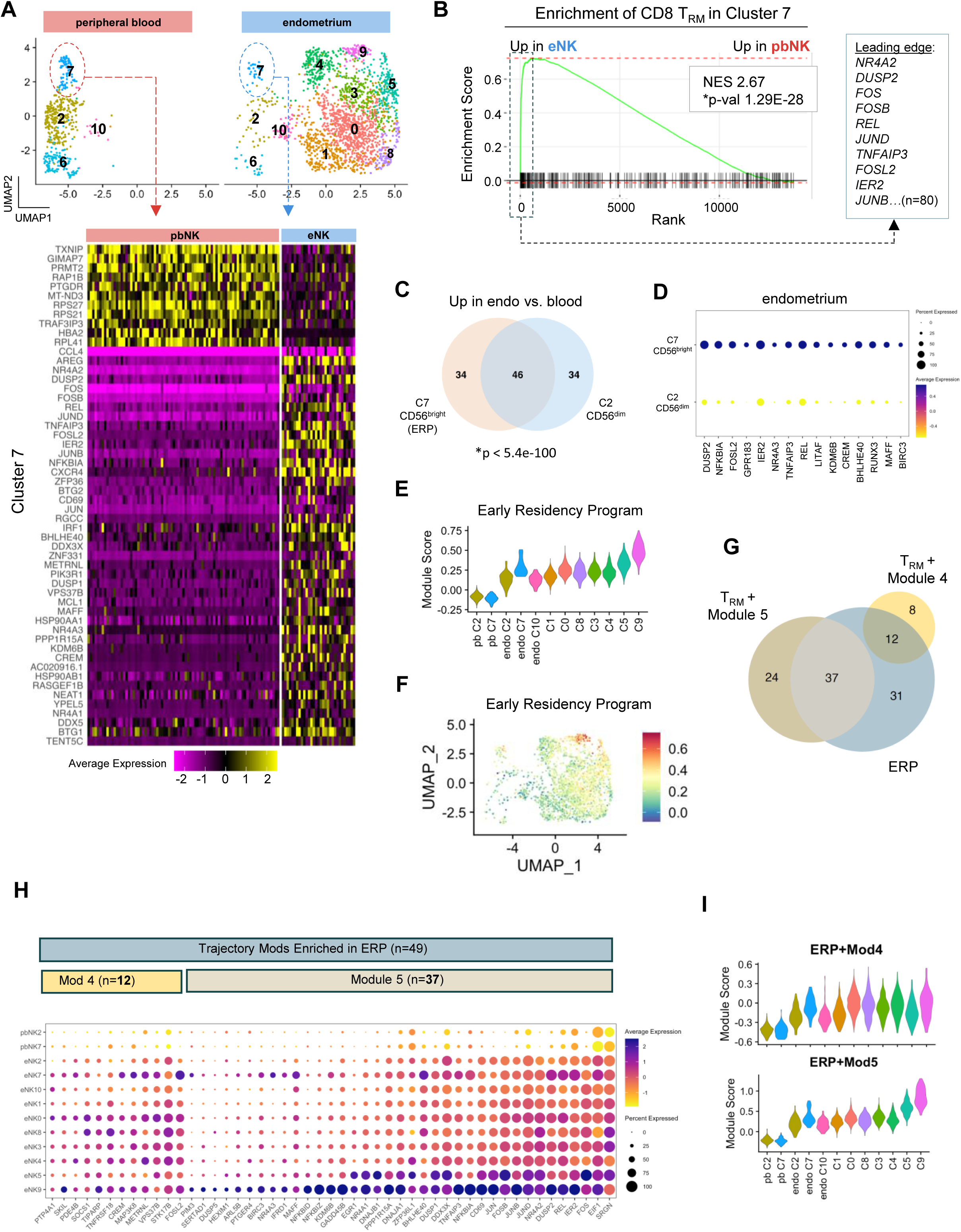
A conserved transcriptional program of early tissue residency distinguishes recent endometrial immigrant NK cells from peripheral blood NK counterparts. (A-I) scRNA-seq results of 2,495 natural killer cells re-clustered from endometrial and peripheral blood CD45+ immune cells. Samples were matched from one individual and collected within 24h during the secretory phase of the menstrual cycle. eNK = endometrial NK; pbNK = peripheral blood NK. (A) Heat map of top differentially expressed genes between pbNK and eNK cells within CD56^bright^ cluster 7. (B) Enrichment of CD8 T_RM_ gene set in genes differentially upregulated in endometrial NK cells versus peripheral blood. NES = normalized enrichment score. Box (hashed) indicates top leading edge genes. (C) Overlap of the ERP signature (n=80 genes) with genes upregulated in endometrial cluster 2 CD56^dim^ cells vs. peripheral blood cluster 2 CD56^dim^ cells. (D) Expression of ERP leading edge genes enriched in endometrial cluster 7 CD56^bright^ cells versus cluster 2 CD56^dim^ cells. (E-F) Violin plot (E) and UMAP rendering of ERP gene program expression across endometrial NK clusters. (G) Overlap of the ERP signature (n=80 genes) with Module 4 genes and Module 5 genes which are upregulated in endometrial trNK across the differentiation trajectory. (H) Expression of individual ERP genes that were shared with trajectory Module 4 or trajectory Module 5. (I) Increasing expression of 12 shared genes (ERP + Mod 4) or 37 shared genes (ERP + Mod 5) across peripheral blood and endometrial clusters.

Although we favored *ITGA1*^lo^*FCGR3A*^lo^ cluster 7 eNK cells as our principal founder NK population, we considered the possibility that *ITGA1*^lo^*FCGR3A*^hi^ cluster 2 eNK cells might also represent a founder population. To discern whether these alternative *ITGA1*^lo^*FCGR3A*^hi^ founder NK cells shared this early CD8 T_RM_ gene signature, we tested for enrichment of the CD8 T_RM_ signature in genes upregulated in C2 eNK cells versus C2 pbNKs. The CD8 T_RM_ signature was highly enriched in C2 eNK cells (Figure S4A), and there was a high degree of overlap between leading edge genes enriched in either C7 eNK cells or C2 eNK cells versus peripheral blood precursors (Figure 4C). Despite the conservation of the ERP between C2 eNKs and C7 eNKs, formal comparison of C7 eNKs vs. C2 eNKs revealed that endometrial founder NK cells potentially derived from CD56^bright^ pbNKs (i.e., C7 eNKs) had higher expression of the ERP (Figures 4D and S4B). We validated these trends in ERP expression in additional samples and datasets and found that the ERP was consistently more highly expressed in eNKs versus their peripheral blood precursors (either CD56^dim^ or CD56^bright^) (Figure S4C). We also found that the ERP was enriched in genes upregulated in CD49a-CD16-compared to CD49a-CD16+ cells in our CITE-seq dataset (Figure S4D), further confirming the increased ERP expression in endometrial *ITGA1^lo^FCGR3A^lo^*founder cells vs *ITGA1^lo^FCGR3A^hi^* founder cells. Moreover, we found enrichment of the CD8 T_RM_ signature among genes differentially upregulated in endometrial NKT cells compared to peripheral blood counterparts (NES 2.69; p-val < 9.1E-25), suggesting that the ERP was a conserved gene program upregulated in newly arrived lymphocytes in different tissues in mice and humans.

Having identified the ERP as a gene program which was differentially expressed between founder NK cells and peripheral blood NK cells, we next tested the hypothesis that ERP genes were among the gene programs promoting the differentiation of trNKs from founder NK cells into transitional states. Consistent with this hypothesis, we found that ERP expression varied across the eNK differentiation trajectory (Figures 4E and 4F). Given that programming of tissue residency was encompassed in gene modules upregulated from founder NK cells through mature populations (Figures 3K and S3C), we defined which genes of the CD8 T_RM_ reference signature overlapped with gene modules with increasing expression from founder NK cells to mature dNK-like cells (Figure S4E). We found that 22% of the CD8 T_RM_ reference gene signature was included in trajectory modules 4 and 5 (Figure S4E), where it comprised 12% and 39% of the module 4 and module 5 signatures, respectively. We next determined that residency genes that increased across the differentiation trajectory in modules 4 and 5 overlapped with the ERP (Figure 4G). Additional examination of ERP gene expression within the module 4 and module 5 gene sets across uNK clusters revealed two distinct expression patterns. We noted that ERP genes shared with module 4 increased in early transitional clusters and then had similar expression between transitional clusters and mature dNK-like clusters (Figures 4H-I and S4F). In contrast, ERP genes shared with module 5 increased between transitional states and mature dNK2-like and mature dNK1-like clusters (Figures 4H-I and S4G). Altogether, these data suggest that a gene program associated with early residency is upregulated in recent NK cell immigrants and changes throughout the differentiation of trNK subsets.

### ERP expression variability underpins endometrial trNK diversity and contributes to transcriptionally divergent mature trNK subsets

As the ERP varied across the differentiation trajectory from founder NK cells to mature dNK-like subsets, we next asked whether the ERP contributed to transcriptional variability between mature dNK-like cells. As our prior analyses were conducted on a limited number of samples, we expanded our analysis cohort and performed scRNA-seq on CD45+ immune cells enriched from secretory phase endometrial biopsies of five additional healthy control volunteers. After aggregation of the samples and selection of NK cells for further analysis, we identified three major groups of NK cells. As before, variability in *ITGA1* expression separated *ITGA1*+ trNK subsets from *ITGA1^lo^* founder NK subsets (Figures 5A and 5B). Given this larger analysis of 14,349 eNK cells from 6 individuals, we could identify a cluster of proliferating cells based on expression of genes associated with cell cycling (i.e., *MKI67*) (Figure 5B), consistent with IL-15-driven proliferation during the secretory phase of the menstrual cycle. Next, we identified dNK-like mature subsets among *ITGA1*+ trNKs after exclusion of low abundance clusters which were predominantly contributed by one sample. After calculation of dNK reference signature performance on these aggregated data, we set thresholds for each signature and made cluster assignments based on our prior criteria. We found two clusters of dNK3-like cells, one cluster of dNK1-like cells, and one cluster of dNK2-like cells (Figure 5C). Approximately half the cells in the dataset could not be assigned to mature dNK-like states (Figure S5A), with unassigned cells comprising the majority of *ITGA1^lo^*founder NK clusters, cycling cells, and a heterogeneous cluster of non-cycling cells that encompassed transitional states (T_0_) (Figure S5A). The identification of the T_0_ transitional cluster among *ITGA1*+ trNK cells was aided by the mapping of barcodes from our prior analysis of matched peripheral blood and endometrium (Figure S5B), where more nuanced transitional states could be discriminated from one another given the presence of circulating pbNKs in the dataset.

**Figure 5.**
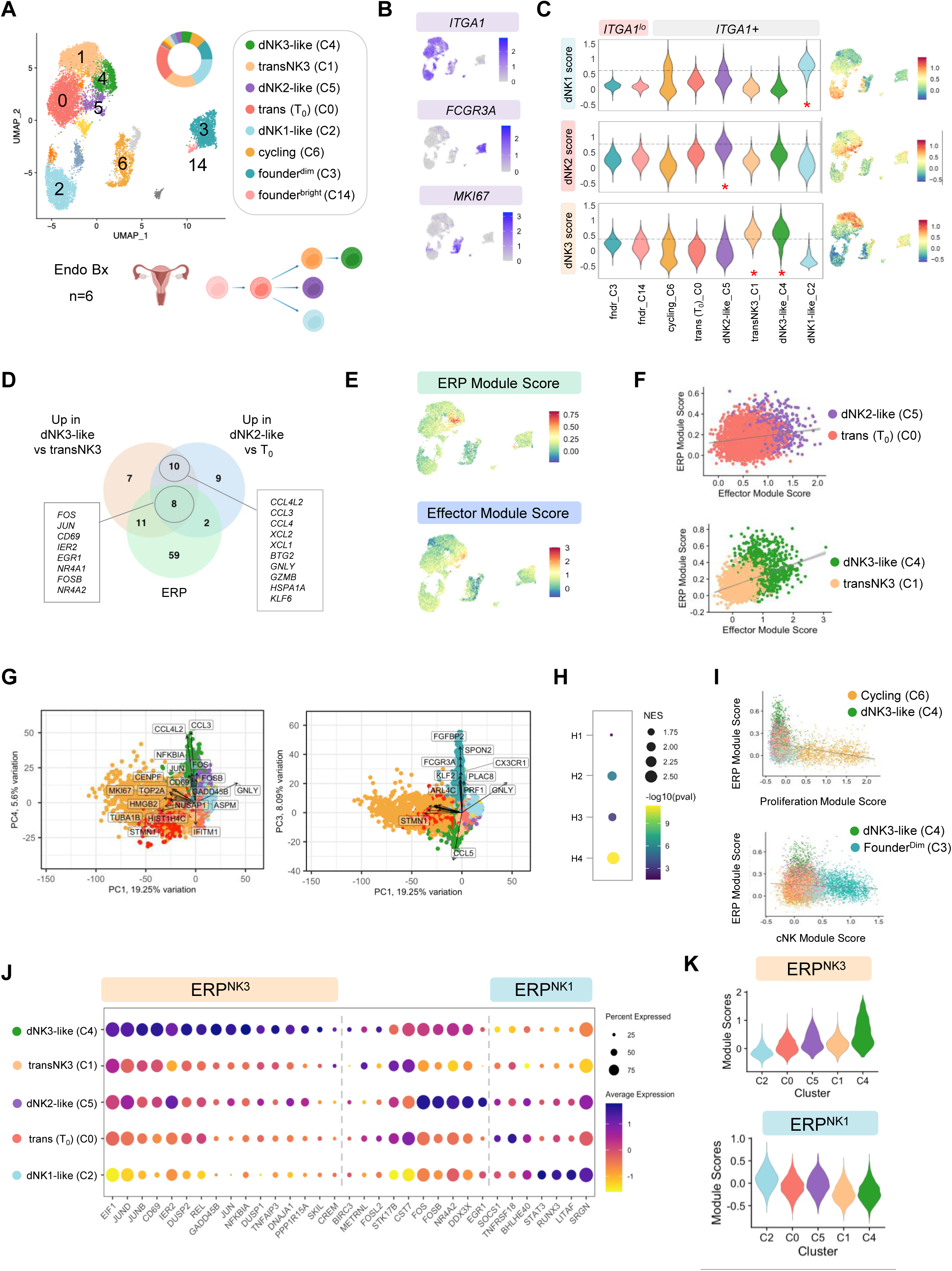
The early residency program (ERP) underpins uterine NK diversity. (A-K) scRNA-seq of 14,349 natural killer cells re-clustered from 41,722 CD45+ immune cells enriched from secretory phase endometrium of six healthy control volunteers. Libraries were aggregated for analysis. (A-C) Identification and enumeration of eNK states. UMAP projection of endometrial NK cells (A, top). Note that clusters of contaminating T cells are colored light gray. Other minor clusters which are not shared across biopsy samples are not annotated. Proposed model of uNK differentiation (A, bottom). Expression of *ITGA1*, *MKI67*, *FCGR3A* among uterine NK cells highlights major transcriptional differences (B). Expression of decidual NK signatures identifies mature endometrial NK clusters (C). (D-E) Co-expression of the ERP with a CCL-based effector gene module in mature dNK2-like and dNK3-like subsets. Venn diagram of overlap between the ERP and differentially expressed genes between dNK2-like cells vs. transitional cells and dNK3-like cells vs. transitional NK3 cells (D). Expression of ERP and CCL-based effector gene module in uterine NK cells (E). Scatter plot of ERP and effector gene module scores demonstrates correlation between ERP expression and the expression of chemokine ligands. (G-I) ERP, cytotoxicity and proliferation gene programs are the most variable transcriptional programs in endometrial NK cells. Biplot representations of PC1 vs. PC4 (left) and PC1 vs. PC3 (right) showcase the contribution of ERP genes to the top PCs in the dataset as well as the relationship of the ERP with genes of the proliferation program (*MKI67*, *CENPF*, etc.) and the cytotoxicity program (*FCGR3A*, *SPON2*) (G). Colors represent individual clusters as in (A), with exception of cycling T cells in red. ERP is enriched in the top four Harmony Embeddings (H1-H4) (i.e., principal components) when genes are ordered within a PC based on their eigenvalue (H). Alternative visualization of negative correlation of ERP with cytotoxicity and proliferation gene modules (I). (J-K) Reciprocal expression of ERP gene modules in dNK3-like and dNK1-like clusters.

We noted that cells with the highest expression of the ERP in our matched blood and endometrial NK analysis mapped again to one of two dNK3-like clusters (Figures 4E and 4F; S5B), supporting the concept that the ERP was differentially expressed between mature trNK subsets. To test this idea, we performed differential expression analysis between mature dNK2-like and dNK3-like cells and their putative precursor clusters (T_0_ and transNK3, respectively). We then compared these differentially expressed genes with the ERP (Figure 5D). Consistent with our hypothesis that the ERP influenced mature dNK-like differentiation, we found substantial overlap between the ERP and DEGs between mature dNK-like cells and their putative precursors (Figure 5D). We also found that expression of a variety of effector genes distinguished mature dNK-like cells from their precursors (Figure 5D), and effector gene expression was highly correlated with expression of the ERP (Figures 5E and 5F). Collectively, these data suggest that the ERP promotes the differentiation of trNKs at later states of maturity and associates with acquisition of effector function in mature dNK-like subsets.

Given the high degree of ERP variability among trNK cells, we speculated that ERP genes would feature prominently in the principal components of our analysis. To test this idea, we first examined the top genes in the principal components (i.e., Harmony embeddings) which explained most of the variance in this dataset (Figures 5G and S5C-D). We find that ERP genes (i.e., *JUN, FOS, CD69*) were among the top contributors to the fourth principal component (Figures 5E and S5C-D). While ERP genes were among the very top contributors to the fourth principal component, we noted that ERP genes were among the largest contributors to each of the four top principal components, accounting for 2.4%, 3.7%, 4.15% and 10% of the top four Harmony loadings. To this point, the ERP was statistically significantly enriched in each of the four Harmony loadings when the genes in each loading were ranked by their eigenvalues (Figure 5H). Taken altogether, the ERP accounted for ∼20% of the total variance in the dataset, on par with genes associated with cytotoxicity (∼11% of variance) and proliferation (∼16% of variance). We further noted that genes of the chemokine effector program which were highly correlated with the ERP (Figures 5D-F) had eigenvectors that were negatively correlated with cytotoxicity genes (Figure 5G). This reciprocal relationship between transcriptional programs of cytotoxicity, proliferation, and the ERP could also be visualized on a scatter plot of the module scores for each program in each cell, where clusters with cells with more expression of the ERP were low expressers of genes associated with cytotoxicity or vice versa (Figure 5I). Altogether, these data indicate that the ERP is a major source of transcriptional variation in endometrial NK cells.

While ERP expression increased progressively from founder NK cells through transitional and into mature dNK2-like and dNK3-like subsets (Figures 4E-F), we noted that expression of the ERP was diminished in dNK1-like cells versus dNK3-like cells (Figures 5E and S5E). To explain this variability among mature dNK-like cells, we hypothesized that dNK1-like cells were downregulating the ERP as they differentiated from transitional subsets. To test this hypothesis, we examined expression of individual ERP genes between dNK1-like and dNK3-like cells. Instead of global downregulation of all ERP genes, we found evidence for modulation of two distinct ERP gene networks by mature dNK-like cells. Although dNK1-like cells had lower expression of AP-1 transcription factor (i.e., *JUN, JUNB, JUND*) as well as NFκB pathway genes (i.e., *REL, NFKBIA*) compared to dNK3-like cells, dNK1-like cells had increased expression of *RUNX3*, *SRGN*, and *STAT3* (Figures 5J and K). Altogether, these results suggest that expression of distinct ERP gene modules promotes the development of two mature trNK subsets in human endometrium.

### Differential sensitivity to TGF-β promotes divergent ERP expression and effector function in endometrial trNK subsets

To gain insight into the tissue factors which might influence ERP expression, we performed a pathway analysis of the ERP genes using Metascape^58^. Pathway analysis suggested that a variety of potential cytokines and growth factors might influence ERP expression through various cellular surface receptors (Figure 6A). From this list of candidates, we elected to focus our subsequent studies on TGF-β for several reasons. First, TGF-β has a well-established role in programming tissue residency in lymphocytes,^30,46,47,49^ and intrinsic TGF-β is required for the development and maintenance of tissue-resident uNKs in mice.^29,30^ Second, TGF-β is expressed in the endometrium, ^35^,^59^and peripheral blood NK cells that have been exposed to TGF-β adopt a decidual NK-like phenotype in vitro.^60^ Although TGF-β is known to upregulate CD103 in peripheral blood NK cells, the ability of TGF-β to upregulate the ERP in pbNK cells has not been tested to our knowledge. To determine whether TGF-β would induce ERP upregulation in pbNK cells, we incubated pbNK cells with or without TGF-β and evaluated *JUNB* and *FOS* mRNA expression using qPCR. We found upregulation of both these genes in cultures containing TGF-β (Figure 6B), illustrating that ERP genes could be upregulated in pbNKs in the presence of TGF-β.

**Figure 6.**
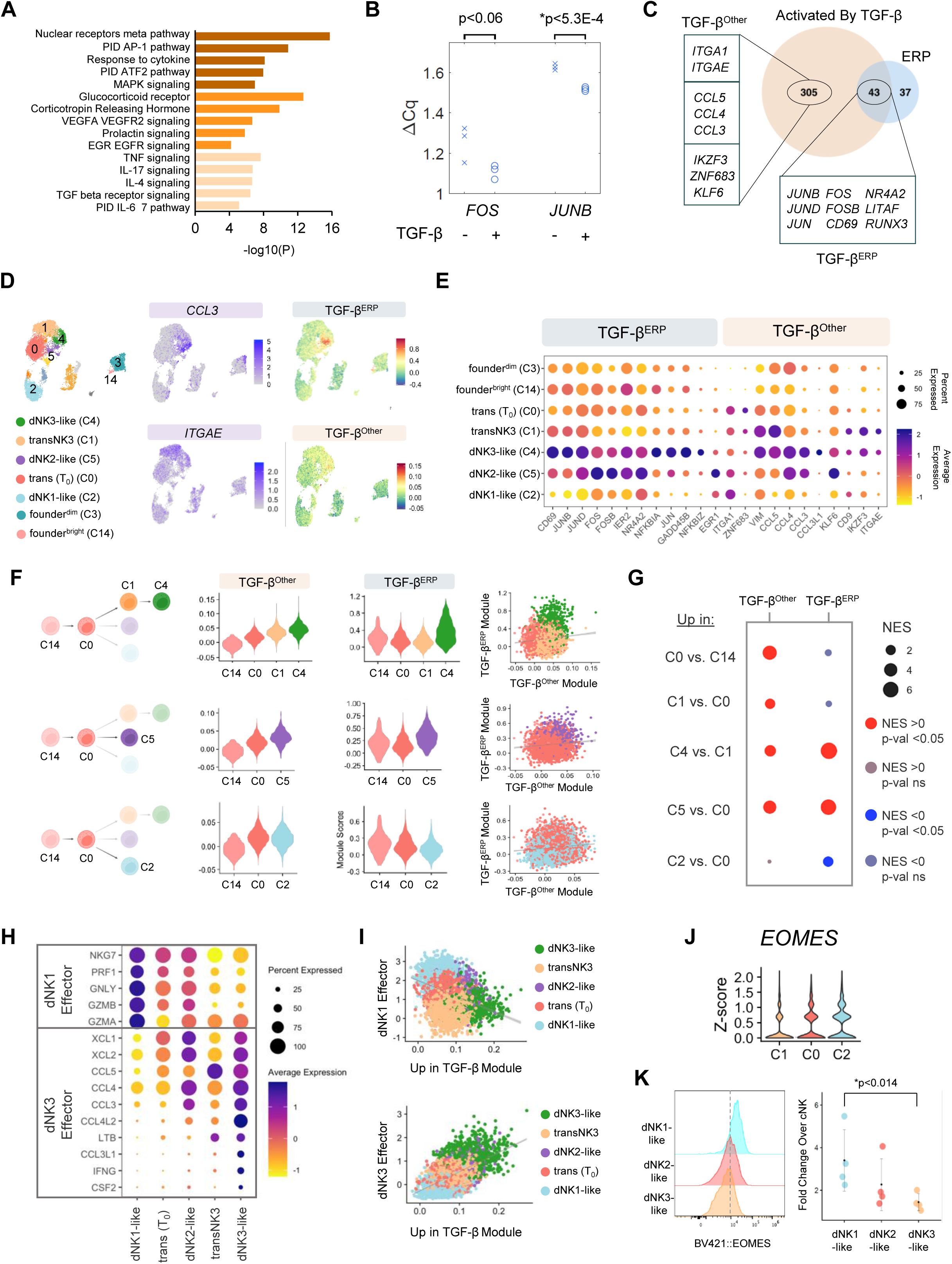
TGF-β modulation of the ERP promotes divergent effector function in eNKs. (A) Pathway analysis reveals enrichment of ERP genes in TGF-β receptor signaling. (B) Upregulation of selected ERP genes (*FOS, JUNB*) in peripheral blood NK cells incubated in vitro with TGF-β for 12 hours. (C) Overlap of ERP with curated list of genes upregulated by TGF-β. (D-J) scRNA-seq of 14,349 natural killer cells re-clustered from 41,722 CD45+ immune cells enriched from secretory phase endometrium of six healthy control volunteers. Libraries were aggregated for analysis. See also Figure 5. (D) Expression of TGF-β^ERP^ correlates with expression of *CCL3*, whereas other genes upregulated by TGF-β (i.e., *ITGAE*) are expressed more diffusely across transNK3 (C1) and dNK3-like (C4) clusters. (E-F) Expression of TGF-β gene targets and gene modules across eNK states. (G) Enrichment of TGF-β gene modules (i.e., TGF-β^Other^ and TGF-β^ERP^) in genes that are differentially upregulated between eNK maturation states. (H-I) Reciprocal expression of effector molecules either upregulated by TGF-β or repressed by TGF-β in dNK3-like clusters or dNK1-like clusters, respectively. (J-K) Eomes expression in eNKs. Eomes is more highly expressed in dNK1-like cells compared to other eNKs. Gene expression in aggregated NK cells from 6 healthy controls (J). Eomes protein expression among eNKs in four additional secretory phase biopsies (K).

Having provided proof of concept that TGF-β could promote the expression of core ERP genes in pbNKs, we next asked whether other ERP genes were targets of TGF-β. We compared the list of 80 ERP genes to a manually curated list of 348 genes upregulated by TGF-β and found that 54% of ERP genes could be activated by TGF-β (Figure 6C). Among the genes activated by TGF-β that were not part of the ERP, we found several integrins of interest (*ITGA1*, *ITGAE*) as well as chemokine ligands (*CCL3*, *CCL4*, *CCL5*) and transcription factors (*ZNF683*, *KLF6*, *IKZF3*) were all expressed to varying degrees among trNK subsets in our data. As many of these ERP genes could be activated by other cytokines, we looked for correlative evidence of TGF-β upregulation of ERP in human endometrial NK cells as we expected that cells with high expression of ERP genes driven by TGF-β would also co-express other TGF-β targets. We thus calculated a module score for expression of TGF-β^ERP^ (n=43 genes) and other genes upregulated by TGF-β (n=305 genes; TGF-β^Other^) in our aggregated data from six healthy control volunteers. We found that clusters of eNK cells with high expression of one TGF-β gene set generally had elevated expression of the other TGF-β gene module (Figure 6D). However, while co-expression of TGF-β gene modules was most apparent in *ITGAE*-expressing dNK3-like cells (Figure 6D), we noted that there was not a 1:1 correlation of the two TGF-β module scores even amongst cells that were the most TGF-β responsive (i.e., *ITGAE*-expressing subsets). These results thus suggested that while TGF-β may be driving ERP expression in these NK subsets, there was heterogeneity in terms of TGF-β responses across the subsets.

To further evaluate patterns of TGF-β responses during uNK development, we examined the expression of individual genes within the TGF-β modules (Figure 6E) as well as overall module expression (Figure 6F) in various uNK developmental states. We also tested for enrichment of TGF-β gene modules among genes upregulated or downregulated across uNK development (Figure 6G). As predicted by our prior analysis (Figures 5E and 5F; Figure 6D), we found that increases in TGF-β^ERP^ expression most correlated with expression of chemokine ligand genes by mature dNK2-like and dNK3-like effector cells (Figure 6E). While increased expression of integrin genes marked the transition from founder to transitional (i.e., *ITGA1*) and ultimately to mature dNK3-like cells for some uNK cells (i.e., *ITGA1*+*ITGAE*+*CD9*) (Figures 6E-F), increases in the expression of AP-1 transcription factors or regulators of NFKB signaling did not track with TGF-β-mediated upregulation of integrins. Instead, increases in *JUN* and *FOS* expression (i.e., TGF-β^ERP^) paralleled upregulation of chemokine ligand genes among more differentiated dNK2-like and dNK3-like effector cells (Figure 6E). Notably, TGF-β responses were more muted in dNK1-like cells (Figures 6F and S6A), as these cells lacked TGF-β-induced upregulation of *ITGAE* or chemokine ligand effector genes (Figure 6E). Notably, while the TGF-β^Other^ module score did not diminish in dNK1-like cells compared to T_0_ transitional cells due to comparable expression of *ITGA1*, *VIM,* and *CD9* and other genes, dNK1-like cells had reduced expression of chemokine ligand genes in concert with reduced expression of the *JUN*, *JUNB*, *JUND* and NFκB elements of the TGF-β^ERP^ module (Figures 6E and S6B). Notably, these specific TGF-β^ERP^ genes were also reduced compared to either *ITGA1^lo^* founder NK population, suggesting active repression during maturation of dNK1-like cells despite the upregulation of *ITGA1*.

Although these analyses suggested that TGF-β was progressively driving trNK development across developmental states, the differences in TGF-β responsiveness between mature dNK-like cells suggested attenuation of TGF-β signaling in dNK1-like cells. To gain insight into the molecular regulators modulating TGF-β responses between mature dNK-like cells in human endometrium, we compared dNK1-like cells with dNK3-like cells and found that cytotoxic effector genes (*GZMB*, *GNLY*) were among the most differentially upregulated genes in dNK1-like cells (Figures S6C-D). Notably, other genes that were differentially expressed in dNK1-like cells vs. dNK3-like cells were transcription factors known to be repressed by TGF-β (Figure S6D), giving additional support to the idea that dNK1-like cells are resistant to TGF-β. As cytotoxic programs are downregulated by TGF-β^61,63^, we examined the correlation between expression of cytotoxic effector genes and expression of TGF-β targets in endometrial NK cells subsets. Consistent with literature in other immune populations, reciprocal expression of cytotoxic gene programs and TGF-β target genes was observed as a continuum across dNK1-like and dNK3-like cells, with transitional trNK cells situated between these two mature subsets (Figures 6H and 6I). Next, we tested the hypothesis that transcriptional regulators of cytotoxic gene programs would differ between dNK1-like and dNK3-like cells. As cytotoxic gene programs are regulated by Eomes^65,67^ and Eomes is known to repress TGF-β-mediated gene programs^46^, we examined Eomes gene and protein expression in trNK subsets. Consistent with its known role as an antagonist of TGF-β programs, Eomes was more highly expressed in dNK1-like cells compared to other populations (Figures 6J and 6K). Altogether, these results suggest that TGF-β induces ERP expression in differentiating tissue resident uNKs and that upregulation of this molecular circuit is linked to expression of immunoregulatory effector molecules.

## DISCUSSION

The goal of this work was to improve our understanding of mechanisms of tissue-resident natural killer cell development in the human endometrium. Using a series of secretory phase endometrial biopsies from healthy control volunteers, we determined that uterine natural killer cells differentiate from a heterogeneous pool of CD49a-negative endometrial immigrants into two dominant CD49a+ populations with divergent effector functions. This transition occurs through a series of developmental intermediates after immigrant cells arrive in the tissue and are stimulated by TGF-β to upregulate a network of transcription factors including AP-1 family members and NR4A2 nuclear receptors. This transcriptional program of early residency, the ERP, is a major molecular contributor to the diversity of uNK cells, as it varied between both founder and mature eNKs as well as between mature eNK subsets. ERP expression is closely linked to immunoregulatory effector functions in dNK3-like mature eNKs and is downregulated in dNK1-like cells with cytotoxic potential. Altogether, these data suggest that the ERP is a developmental circuit induced by TGF-β upon tissue entry that guides the acquisition of effector function in differentiating uterine natural killer cells.

A critical step in the creation of this developmental framework was the identification of the uterine NK founder population. Our comprehensive multimodal assessment of the endometrium revealed that uNK founder cells are a heterogeneous population that share the phenotype of their CD56^bright^ and CD56^dim^ peripheral blood precursors, thus explaining why these cells have historically been interpreted as blood contaminants. Moreover, the earliest transcriptional changes associated with tissue entry and the founder NK state are subtle, making it challenging to discern these changes in a highly dynamic tissue using noisy single-cell datasets. We were able to discriminate founder NK cells from their blood-based precursors because of several intentional features of our experimental design. These features included direct linkage of cellular transcriptomes with surface phenotypes using CITE-seq, and comparison of matched peripheral blood samples with endometrial biopsies at consistent and targeted time points in the menstrual cycle.

The identification of founder cells and subsequent developmental intermediates was further facilitated by our novel approach to NK cell state assignments based on reference gene set expression thresholds, allowing the assignment of multiple states. This latter feature proved invaluable as it enabled the identification of cells sharing one or more transcriptional signatures which is key to identifying cells in transition from one transcriptional state to another. We were additionally aided by the application of knowledge gained from studies of tissue resident lymphocytes in mice and other human organs. In particular, the recent discovery that CD8 T_RM_ upregulate immediate early genes prior to expression of integrins during early differentiation in mucosal tissues^45^ allowed us to identify the putative founder NK populations and the subsequent developmental states in human tissues where complementary lymphocyte tracking and time course studies cannot be performed.

Our identification of the early residency program in human NK cells advances our understanding of tissue resident biology in several important ways. First, it underscores the importance of this network of transcription factors in lymphocyte biology. Despite evidence through genetic knockdown experiments that immediate early genes (IEGs) such as *JUN* and *FOS* mediate critical aspects of tissue residency^45^, some investigators have reported exclusion of these genes in their analysis of tissue resident NK cells^56^, indicating uncertainty about the relevance of this gene program. However, we have observed in prior work that reductions in expression of AP-1 transcription factors and nuclear receptors in endometrial NK cells associated with loss of tissue resident cells and adverse pregnancy outcomes in human transplant recipients.^14^ The current description of the ERP as a transcriptional program governing the development of tissue resident NK cells in humans thus synergizes with our prior work and supports the seminal findings of Kurd et al.^45^ that this molecular circuit has a key role in the development of tissue residency.

Another important contribution of our study is the identification of the ERP in innate lymphocytes that do not express an antigen receptor. As immediate early genes that were previously described in developing CD8 TRM are often induced by antigen receptor signals, it is reasonable to conclude that antigen plays a role in the programming of tissue residency in T cells. Although there may be many roles for antigen and antigen receptor signaling in the programming of T_RM_, our data provide strong evidence that other cellular receptors, likely cytokine receptors, also play a role in tissue residency programming.

How lymphocytes integrate cytokine receptor signals and/or antigen receptor signals during tissue-resident lymphocyte development is an important area of future study. Moreover, while we attribute the modulation of ERP expression during trNK development to TGF-β in our work, additional investigations will be required to characterize the suite of cytokines which may activate or repress the early residency program.

We utilized changes in expression of the early residency program from the blood to mature trNKs to determine that many endometrial NK cells exist in a state of transition as developmental intermediates. Our phenotypic and transcriptional studies of these developmental intermediates further suggest that these cells largely correspond to the dNK2 population proposed by Vento-Tormo et al^17^. However, whether developmental intermediates we have identified in the endometrium exist in the decidua is unclear, and additional studies will be needed to determine to what extent the decidualizing microenvironment and/or the presence of trophoblast during pregnancy impact these transitions and uterine NK development overall.

The developmental framework we propose with this work relies on our ability to identify starting points of differentiation. While we favor a developmental model where mature NK cells from the peripheral blood seed the endometrium, our model does not exclude the contribution of potential earlier precursors such as ILCPs to the tissue-resident NK pool. Whether the ERP plays a role in differentiation from alternative precursors will be an important area of future investigation, and these experiments will critically synergize with additional studies to better define the contribution of peripheral blood CD56^bright^ and CD56^dim^ cells to the founder pool as well. Given that the ERP was more highly expressed in endometrial NK cells co-expressing the CD56^bright^ signature than the CD56^dim^ signature, we favored a reductionist model whereby founder NK cells derived from the peripheral blood CD56^bright^ pool provide the primary input into the precursor pool. Nevertheless, our data suggest that CD56^dim^ cells can be significantly altered upon tissue entry, consistent with the work of others who have described the spectrum of effects of TGF-β on peripheral blood CD56^dim^ cells.^60,62^ Moreover, recent studies in mice continue to improve our understanding of the diverse cell types – including mature cNKs - which traffic into tissues during homeostasis and infection and become tissue-resident NK cells^32,64^. We anticipate that our understanding of the role of the ERP in lymphocyte residency will continue to evolve as we improve our understanding of the founder NK pool.

Our interpretation of the role of the ERP in tissue resident NK development also depends on the endpoint of differentiation we select for our developmental model. We chose to identify endometrial NK developmental endpoints based on expression of reference gene signatures derived from transcriptional differences between decidual NK cell subsets^17^. The cells we identified as endpoints in our developmental model (i.e., dNK1-like and dNK3-like cells) expressed effector programs defined by chemokine ligands and cytotoxic molecules, respectively. Although the use of reference gene signatures is a common way of annotating cell types, the genes included in such lists are somewhat arbitrary in nature, subject to bias, and are highly dependent on the method of derivation. Moreover, there are currently no agreed upon standards or external benchmarks to reproducibly assign or to otherwise verify a cell type or cell state for uterine NK cells. The development of functional assays for NK cells of reproductive tissue will aid in assessing cellular assignments and the presence of differentiation endpoints.

Related to functional states, we note that the co-expression of the ERP with these effector molecules is consistent with the role that the AP-1 complex and NF-κB family play in activating effector function. We are especially intrigued by the possible role of the ERP in repression of cytotoxic function, which our data suggest corresponds to downregulation of Eomes by TGF-β in maturing dNK3-like cells. Equally intriguing is the mechanism by which TGF-β and the ERP is potentially antagonized in developing dNK1-like cells. Our data point to a molecular circuit involving Eomes and the Eomes regulon. However, whether Eomes is the primary determinant of dNK1-like cell modulation of TGF-β responses or merely an indirect consequence of some other upstream regulator remains to be determined. Indeed, the mechanisms by which dNK1-like cells maintain tissue residency in the face of TGF-β antagonism are unclear at this juncture, but we are intrigued by recent evidence that molecular regulators which convey cytotoxic capacity (i.e., RUNX3, EOMES) may also function to promote tissue residency.^44,66,68^ Additional work is needed to understand the specific cytokines and environmental signals which may support the differentiation of tissue-resident populations that maintain cytotoxic capacity.

In conclusion, we identify a molecular circuit governing human tissue-resident uNK development that is conserved across species, tissues, and cell types. This circuit is modulated at different developmental stages and appears closely tied to the acquisition of immunoregulatory function. ERP genes are induced by TGF-β and the degree of ERP expression tracks with TGF-β responses across the developmental stages, suggesting that a core function of the ERP is to mediate TGF-β signals. As ERP genes overlap with previously described immediate early response genes and are targets of diverse cytokines, we speculate that the ERP serves as an integration circuit that synthesizes TGF-β signals with other cytokines to prepare tissue immigrant lymphocytes for new functionality. Collectively, these data improve our understanding of the molecular mechanisms of tissue resident lymphocyte development and may prove to be of benefit as we discover how perturbations of tissue resident lymphocytes either contribute to disease pathogenesis or can be intentionally engineered for therapeutic purposes.

## ACKNOWLEDGEMENTS

Stefani D. Yates, Jayme E. Locke, Sunita Patel, Gaby Halder, Shanrun Liu, Vidya Sagar Hanumanthu.

## RESOURCE AVAILABILITY

### Lead contact

Further information and requests for resources and reagents should be directed to the lead contact, Dr. Paige Porrett.

### Materials availability

No unique reagents were generated during the course of this study.

### Data and code availability

All sequencing data that support the findings of this study have been deposited in the NCBI Gene Expression Omnibus (GEO) under accession numbers GSE292828 and GSE292829 for scRNA-seq and CITE-seq data, respectively. All sequence analyses were performed using open-source software and packages. Signature score thresholding was performed in MATLAB. Details of the code functions used from respective packages are provided in the Methods section. Generated code and all corresponding analytic details are available on GitHub at: https://github.com/PorrettLab/A-developmental-program-of-early-residency-promotes-the-differentiation-of-divergent-uNK-cells/tree/main and on Zenodo at: https://doi.org/10.5281/zenodo.15725787. Any additional information required to reanalyze the data in this paper is available upon request from the lead contact.

## RESOURCE AVAILABILITY

### Materials availability

No unique reagents were generated during the course of this study.

## STAR★Methods

Detailed methods are provided in the online version of this paper and include the following:

### KEY RESOURCES TABLE

#### EXPERIMENTAL MODEL AND STUDY PARTICIPANT DETAILS

##### Human

###### Ethics Approval and Participant Consent

The study titled “Mechanisms of Uterine NK Cell Differentiation” received ethics approval from the University of Alabama at Birmingham Institutional Review Board (IRB-300006859). Prior to participation, all participants were required to provide written informed consent.

###### Participant Recruitment and Screening

Eligible participants were premenopausal women aged 18-50, who were deemed suitable for the study based on a detailed screening questionnaire that included gender, race, ethnicity, and obstetric and gynecological history. Ancestry and socioeconomic status were not collected as part of this questionnaire, as the lack of this information does not limit the generalizability of this study. The primary aim of this questionnaire was to assess each participant’s compatibility with the study requirements. Exclusion criteria were rigorously applied to ensure the selection of a homogeneous participant pool devoid of confounding variables to the extent possible. These criteria included the absence of a history of malignancy, uncontrolled diabetes, pharmacologic immune modulation, systemic autoimmune disease, prior organ or bone marrow transplantation, chemotherapy within the preceding three years, pelvic radiation treatment, chronic or end-stage kidney disease (defined as dialysis dependence or a Glomerular Filtration Rate (GFR) < 60), chronic or end-stage liver disease, HIV infection, recent instrumentation of the uterine cavity (e.g., Dilation and Curettage (D&C)) within the last 12 months, use of an Intrauterine Device (IUD) within the past 3 months, oral contraceptive use within the past 3 months, pregnancy within the preceding 12 months, NSAID or aspirin use within the previous 10 days, or any medical contraindication to NSAID usage. Additional exclusion criteria targeted women currently using oral contraceptive agents, anti-coagulants, aspirin, those who were pregnant, had an indwelling IUD, or were infected with Hepatitis C Virus (HCV) or Hepatitis B Virus (HBV).

#### Healthy Control Biopsy Collection

The study utilized uterine biopsies and blood from consenting women aged between 27 and 46 years, inclusive of diverse ethnic backgrounds, including individuals who self-identified as Caucasian, African American, and Asian. This approach ensured a representative sample that could provide insights into the mechanisms of uterine NK cell differentiation across a broad demographic spectrum.

Participants were screened for eligibility based on the criteria prior to their clinic visits at the University of Alabama at Birmingham. Eligible participants were then approached for consent, ensuring informed participation in the study.

#### Healthy Control PBMC Collection

The study utilized blood collected from consenting women who had met the criteria for the healthy control uterine biopsy collection. This approach ensured a representative sample that could provide insights into the similarities and differences between peripheral blood NK cells and uterine NK cells. Blood was also collected to isolate NK cells for *in vitro* incubation with TGF-β.

## METHOD DETAILS

### Endometrial Biopsy and Single-Cell Isolation

Endometrial tissue was used for spectral flow cytometry, cellular indexing of transcriptomes and epitopes sequencing (CITE-seq), and single-cell RNA sequencing (scRNAseq).

Participants were given Clearblue digital ovulation tests (SPD Swiss Precision Diagnostics GmbH; Item number 245-03-0301) to take home and track their ovulation date, following the manufacturer’s instructions. Once a participant received a solid smile on the test, they would reach out to schedule their endometrial biopsy. These biopsies were scheduled for dates during the secretory phase of the menstrual cycle.

Endometrial biopsies were collected in a clinical setting with a Pipelle suction curette (Cooper Surgical; Ref. 8200) and kept in Dulbecco’s Phosphate Buffered Saline (1X) with Calcium and Magnesium (Gibco; Cat. # 14040-117), hereafter referred to as DPBS+/+, on ice until digestion. The collection tube and media were weighed before and after tissue collection to calculate the tissue weight.

On ice, biopsies were finely minced with a razor blade in a petri dish and then transferred to a 50mL conical tube. Enough pre-warmed 37°C DPBS+/+ was then added to fully cover the tissue. A 1:100 dilution of 5mg/mL Liberase TM Research Grade (Sigma-Aldrich; Cat. # 5401127001) and a 1:1000 dilution of 1mg/mL DNAse I (Sigma-Aldrich; Cat. # 10104159001) were then added to the pre-warmed DPBS+/+ to create the digestion mixture. For example, if 5mL of DPBS+/+ were used to fully cover the tissue, then 40µL of 5mg/mL Liberase TM and 5µL of 1mg/mL DNAse I would be added.

Digestion was performed in an Incu-Shaker H1001-M (Benchmark Scientific) at 37°C and 240 RPM for 10-30 minutes, with checks every 5 minutes to ensure the tissue was neither over- nor under-digested. Once fully digested, the conical was placed on ice and the sample was filtered through a 70µm strainer (Stemcell Technologies; Cat. # 27260) into a fresh 50mL conical tube. The strainer was rinsed three (3) to five (5) times with 1mL ice-cold DPBS+/+. The digested biopsy was then centrifuged at 300g for 5 minutes at 4°C, and the supernatant was aspirated. The cell pellet was incubated in 3mL ACK lysis buffer (Quality Biological; Cat. No. 118-156-101) and transferred to a 15mL conical on ice for 2 minutes. At the end of cell lysis, 12mL ice-cold Dulbecco’s Phosphate Buffered Saline (1X) without Calcium and Magnesium (Gibco; Cat. No. 14190-136), hereafter referred to as DPBS-/-, was added to the cell suspension to wash. The cells were then centrifuged at 300 g for 5 minutes at 4°C. The buffer was removed, and 0.5 – 1mL of fresh cell staining buffer was added to resuspend the cells for counting. The cell staining buffer for samples meant for scRNA sequencing consisted of ice-cold DPBS-/- with 0.04% (w/v) BSA (Jackson ImmunoResearch; Cat. No. 001-000-162), and this buffer was also used for sorting. The cell staining buffer for all other samples consisted of ice-cold DPBS+/+ with 2% (v/v) FBS (Gemini Bio; Ref. 100106) and 0.4% (v/v) 0.5M EDTA (Fisher Scientific; Cat. No. AAJ15694AP).

### PBMC Isolation

PBMCs were isolated from whole blood for spectral flow cytometry, CITE-seq, and scRNAseq.

Blood was collected in EDTA-K2 tubes vacutainers (BD and Company; Ref. 366643) and blood collection set (MYCO Medical Supplies, Inc.; Ref. GSBCS23G-7T). StemCell’s SepMate protocol was followed to isolate PBMCs using Lymproprep (Stemcell Technologies; Ref. 07811) and either a Sepmate-15 or −50 (Stemcell Technologies; Ref. 85420 or 85450), depending on total blood volume.

The only deviation from the StemCell’s protocol was in Step 4. All blood samples were centrifuged for 20 minutes. Additionally, the wash to remove platelets was always performed. After removing the final wash buffer, any remaining RBCs were lysed using 3mL of room-temperature ACK lysis buffer for 2 minutes on ice. Ice-cold DPBS was added to the 14mL mark and mixed to finish the lysis reaction. Cells were pelleted by centrifugation, 400g for 5 minutes at 4°C. The buffer was removed, and 0.5-1mL fresh sort buffer was added to resuspend the cells, which were then counted as described in the “Cell Counting” Methods section. PBMCs were then either used for scRNA library preparation, *in vitro* incubation with TGF-β, or frozen as detailed in the “Cell Freezing and Thawing Methods” section.

#### NK Cell Isolation from Whole Blood

NK cells were isolated from whole blood using the MACSxpress Whole Blood NK Cell Isolation kit, human (Miltenyi Biotech, cat 130-127-695) for *in vitro* incubation with TGF-β.

The blood was incubated with an antibody isolation cocktail for 5 minutes and centrifuged at 50 g for 1 minute. Non-NK leukocytes were depleted by placing the sample in a magnetic field for 15 minutes. The enriched, supernatant cell suspension was collected and centrifuged at 350 g for 10 minutes. To remove residual erythrocytes, the cell pellet was subjected to 10mL of ACK lysis buffer for 5 minutes at room temperature, washed, and centrifuged at 350 g for 10 minutes at 4°C. The final cell pellet was resuspended for cell counting and downstream applications.

Purity was assessed after NK cell isolation by staining whole blood and freshly isolated cells. Refer to the “Live/Dead Staining,” “Surface Staining for *in vitro* Incubation,” “Cell Fixation and Permeabilization,” and “Internal Staining for *in vitro* Incubation” subsections of “Cell Staining” for the protocols used.

#### Cell Freezing and Thawing

Cells that were not immediately used for sequencing were pelleted by centrifugation at 300 – 400 g for 5 minutes at 4°C, and then resuspended in freezing buffer, which consisted of 90% (v/v) FBS supplemented with 10% (v/v) DMSO (Fisher Bioreagents; Ref. BP231-100). Cells were then aliquoted into cryovials and then frozen at −80°C in a Mr. Frosty Freezing Container (Thermo Fisher Scientific; Ref. 15-350-50) for 24 – 48 hours before being placed in liquid nitrogen cryostorage.

Cryopreserved cells were partially thawed in a 37°C water bath while gently swirling. After half the volume of the cells was thawed, 500µL of 37°C-warmed thawing media, RPMI (Gibco; Ref. 11875-093) supplemented with 10% FBS, was added to the cryovials. Suspended cells were transferred to a 15mL conical tube containing 9mL of warmed thawing media. Cells were centrifuged at 300 – 400 g for 5 minutes at 4°C. The thaw media was aspirated, and the cells were washed again in thaw media. After centrifugation, the thaw media was aspirated, and cells were suspended in sorting buffer. Cells were then stained, sorted, and counted as above before being used for scRNA experiments.

### Cell Counting

Cells were counted twice in either a 1:10 or 1:100 dilution of cells to 0.04% Trypan blue in PBS (Gibco; Ref. 15250-061). Cell counts were performed using a DHC-N01-5 disposable Neubauer improved hemocytometer (INCYTO; DHC-N01-5). To reduce human error, 10µL of the diluted cells were added to each well to obtain a mean cell count.

Cell concentration was calculated with the following equation, where DF is the dilution factor:

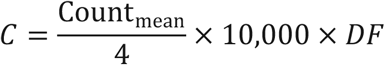

Cell number was calculated with the following equation, where V_res_ is the resuspension volume:

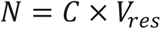

#### *in Vitro* NK cell TGF-β Incubation

All NK cells isolated as described in the NK Cell Isolation from Whole Blood section were incubated in culture media, defined as: RPMI, 10% FBS, and 50µM beta-mercaptoethanol (Gibco; Ref. 21985-023). Cells were plated in 48-well plates (Thermo Fisher Scientific; 12565322). Each well received 1 million cells in 1mL of culture media. Four time points were used for this study: 3, 12, 24, and 72 hours in a 37°C incubator with 5% CO_2_. Experimental wells contained 10 ng/mL Human TGF-β1 (PeproTech Inc.; 100-21-10UG) reconstituted in 10mM Citric Acid, pH 3.0 to 0.1 mg/mL (Fisher Bioreagents; Ref. BP339-500) while the control wells only received UltraPure distilled water. Cells incubated for 72 hours also received 5ng/mL IL-15 (PeproTech Inc.; 200-15-10UG) to ensure cell viability. Each experimental condition was performed in duplicate for each time point, except for the 72-hour time point, which was conducted in quadruplicate to allow for both flow cytometry and RNA extraction for quantitative PCR (qPCR). The duplicates for each time point were pooled for RNA extraction.

#### Cell Staining

##### Live/Dead Staining

All cells to be used for spectral flow cytometry or scRNA sequencing were separated into samples containing 1 million cells or fewer, and each aliquot was washed with 1mL of DPBS+/+. Cells were then centrifuged to pellet at 300 – 400 g for 5 minutes at 4°C. The supernatant was aspirated, and the pellet resuspended in 150µL DPBS+/+ with 0.25µL LIVE/DEAD Fixable Aqua Dead Cell Stain Kit (Invitrogen; Cat. L34957) and incubated in the dark, on ice for 30 minutes. After the incubation, cells were washed with 1mL cell staining buffer, as defined in the “Endometrial Biopsy and Single Cell Isolation” Methods section, and centrifuged to pellet as previously described in this section before proceeding to surface staining.

##### Surface Staining for Sort

After Live/Dead staining, the cells were resuspended in 100µL of sorting surface stain master mix and incubated for 30 minutes in the dark on ice. After incubation, cells were washed with 1mL of cell staining buffer and centrifuged to pellet as described in the “Live/Dead Staining” Methods section.

The sorting surface stain master mix was created using 1µL Alexa Fluor 700 anti-human CD45 antibody (BioLegend Cat. # 368514, RRID:AB_2566374), 1µL PE anti-human CD235a (Glycophorin A) antibody (BioLegend Cat. # 349105, RRID:AB_10641707), and 100µL cell staining buffer per sample.

##### CITE-seq Staining

Cells to be stained for CITE-seq were stained in accordance with BioLegend’s “TotalSeq-B with 10x Feature Barcoding Technology” protocol, starting with Step 3.2.1, and BioLegend’s “TotalSeq Antibody Cocktail “Spike-in” Guidance” protocol.

The reconstitution volume for the TotalSeq™-B Human Universal Cocktail, V1.0 (BioLegend; Cat. # 399904, RRID:AB_2892472) was 17.5µL of either Cell Staining Buffer (Biolegend; Cat. No. 420201) or the DPBS-/- with 0.04% (w/v) BSA cell staining buffer and 10µL of the sorting surface stain master mix. The cells to be stained were resuspended in 22.5µL of cell staining buffer and 2.5µL of Human TruStain FcX Fc Receptor Blocking Solution (BioLegend; Cat. # 422301). This resulted in a total staining volume of 50µL.

The only deviations from BioLegend’s “TotalSeq-B with 10x Feature Barcoding Technology” protocol were in Step 3.2.1, where up to 2 million cells were stained per TotalSeq-B Human Universal Cocktail, and Step 5.1.6, where cells were only washed once instead of three times to both save time and increase cell viability.

##### Surface Staining for Spectral Cytometry

After Live/Dead staining, cells to be analyzed by spectral flow cytometry were resuspended in 100µL of surface stain master mix per sample and incubated in the dark on ice for 30 minutes. After incubation, cells were washed with 1mL of cell staining buffer and centrifuged to pellet as described in the “Live/Dead Staining” Methods section.

The surface stain master mix consisted of the following for each sample: 30µL BD Horizon Brilliant Stain Buffer (BD Biosciences; Cat. # 563794), 1µL APC anti-human CD56 antibody (BioLegend; Cat. # 318310, RRID:AB_604106), 0.5µL APC/Cyanine7 anti-human CD3 antibody (BioLegend; Cat. # 317342, RRID:AB_2563410), 1µL Brilliant Violet 421 anti-human CD103 antibody (BioLegend; Cat. # 350214, RRID:AB_2563514), 2µL Brilliant Violet 711 anti-human CD49a antibody (BD Biosciences; Cat. # 742361, RRID:AB_2740719), 2µL FITC anti-human CD158a/h (KIRS) antibody (Miltenyi Biotec; Cat. # 130-118-973, RRID:AB_2733621), 1µL FITC anti-human CD158b1,b2,j (KIRS) antibody (BD Biosciences; Cat. # 556070, RRID:AB_396339), 1µL FITC anti-human CD158e1 (KIRS) antibody (BioLegend; Cat. # 312706, RRID:AB_314945), 2µL FITC anti-human CD158i antibody (Miltenyi Biotec; Cat. # 130-114-614, RRID:AB_2655362), 1µL Pacific Blue anti-human CD45 antibody (BD Biosciences; Cat. # 556070, RRID:AB_396339), 1µL PE/Cyanine7 anti-human CD39 antibody (BioLegend; Cat. # 328212, RRID:AB_2099950), 1µL PE/Dazzle594 anti-human CD16 antibody (BioLegend; Cat. # 302054, RRID:AB_2563639), 1µL PerCP anti-human CD14 antibody (BioLegend; Cat. # 325632, RRID:AB_2563328), and 56.5µL cell staining buffer.

##### Surface Staining for in vitro Incubation

After Live/Dead staining, cells from the 72-hour incubation with TGF-β were resuspended in 100µL of surface stain master mix per sample and incubated in the dark on ice for 30 minutes. After incubation, cells were washed with 1mL of cell staining buffer and centrifuged to pellet as described in the “Live/Dead Staining” Methods section.

The surface stain master mix consisted of the following for each sample: 20µL BD Horizon Brilliant Stain Buffer, 3µL Alexa Fluor 532 anti-human CD14 antibody (Thermo Fisher Scientific; Cat. # 58-0149-41, RRID:AB_11218093), 2µL APC anti-human CD103 antibody (BioLegend Cat. # 350216, RRID:AB_2563907), 1µL APC/Cyanine7 anti-human CD3 antibody, 1µL Brilliant Violet 570 anti-human CD8a antibody (BioLegend; Cat. # 301038, RRID:AB_2563213), 1µL Brilliant Violet 650 antihuman CD4 antibody (BioLegend; Cat. # 317436, RRID:AB_2563050), 0.5µL FITC anti-human CD158a/h (KIRS) antibody, 0.5µL FITC anti-human CD158b1,b2,j (KIRS) antibody, 0.5µL FITC anti-human CD158e1 (KIRS) antibody, 0.5µL FITC anti-human CD158i antibody, 2µL PE anti-human CD19 antibody (BioLegend; Cat. # 302208, RRID:AB_314238), 3µL PE/Cyanine5 anti-human CD56 antibody (BioLegend; Cat. # 362516, RRID:AB_2564089), 2µL PE/Cyanine7 anti-human CD49a antibody (BioLegend; Cat. # 328312, RRID:AB_2566272), 1µL PE/Dazzle594 anti-human CD16 antibody, and 54µL cell staining buffer.

##### Cell Fixation and Permeabilization

The only cells not fixed and permeabilized were those intended for sorting and single-cell RNA sequencing (scRNA-seq). All other cells were fixed and permeabilized using the eBioscience Foxp3 / Transcription Factor Staining Buffer Set (Thermo Fisher Scientific; Cat. # 00-5523-00).

After surface staining, cells were resuspended in 300µL of Fix/Perm working solution per sample and incubated in the dark on ice for 30 minutes. The working solution was created using a 1:4 ratio of Fixation/Permeabilization Concentrate to Fixation/Permeabilization Diluent.

After incubation, the cells were washed with 1mL of Fix/Perm buffer, which was prepared in a 1:10 ratio of 10X Permeabilization Buffer to UltraPure Distilled Water (Thermo Fisher Scientific; Cat. #10977015). Cells were then centrifuged to pellet and either moved on to internal staining or resuspended in 200µL Fix/Perm buffer for spectral flow cytometry.

##### Internal Staining for in vitro Incubation

After cell fixation and permeabilization, cells from the 72-hour incubation with TGF-β were resuspended in 100µL of internal stain master mix per sample and incubated 30 minutes in the dark at 4°C. After incubation, the cells were washed with 1mL of Fix/Perm buffer, centrifuged to pellet as previously described in the “Live/Dead Staining” Methods section, and resuspended in 200µL Fix/Perm buffer for spectral flow cytometry.

The internal stain master mix consisted of the following for each sample: 20µL BD Horizon Brilliant Stain Buffer, 2µL Brilliant Violet 421 anti-human EOMES antibody (BD Biosciences; Cat. # 567166; RRID:AB_2916483), 2µL Brilliant Violet 785 anti-human T-bet antibody (BioLegend; Cat. # 644835; RRID:AB_2721566), and 76µL Fix/Perm buffer.

### Fluorescence-activated cell sorting (FACS)

Live, CD45+, CD235a-cells were sorted on a BD FACSAria I (RRID:SCR_019595) or BD FACSAria II Cell Sorter (RRID:SCR_018934) at the UAB Flow Cytometry Core Facility into 2mL RNase-/DNase-free microcentrifuge tubes with 50µL of sorting buffer at the bottom. Single-color controls for surface marker fluorophores were created using the AbC Total Antibody Compensation Bead Kit (Thermo Fisher Scientific, Cat. # A10497) and the same antibodies used to stain the cells. Brilliant Violet 510 anti-human CD8a antibody (BioLegend; Cat. # 300934; RRID:AB_2814115) was used to compensate for Live/Dead Aqua. Cells were recounted post-sort to determine cell number, concentration, and viability.

### Spectral Flow Cytometry

Cells stained for spectral flow cytometry were analyzed using a Cytek Northern Lights spectral cytometer (RRID:SCR_027072). Single-color controls for surface and internal fluorophores were created with the AbC Total Antibody Compensation Bead Kit and the same antibodies used to stain the cells. The ArC Amine Reactive Compensation Bead Kit (Thermo Fisher Scientific; Cat. # A10346) and LIVE/DEAD Fixable Aqua Dead Cell Stain were used to create the single-color control for Live/Dead.

### CITE-seq and scRNAseq Library Preparation

Sorted cells, with or without CITE-seq antibody staining, were used for 5’ or 3’ scRNA library preparation. scRNA GEMs and libraries were prepared according to the manufacturer’s instructions. 5’ libraries were prepared according to the 10x Genomics CG000399 protocol, Rev B using the Chromium Next GEM Single Cell 5’ Kit v2 (10x Genomics; PN-1000263). 3’ libraries were prepared using 10x Genomics CG000331, Rev E protocol, or CG000206, Rev D if a CITE-seq antibody panel was used with the appropriate Chromium Next GEM Single Cell 3ʹ Reagent Kit v3.1 (10x Genomics; PN 1000121). CITE-seq libraries were amplified using Chromium GEM-X Single Cell 3 Feature Barcode Kit v4 (10x Genomics; Cat. # 1000702). Both 3’ and 5’ libraries Dual Index Kit TT Set A (10x Genomics; PN 3000431) were used to finalize libraries. The Dual Index Kit NT Set A (10x Genomics; Cat. # 1000242) was used to finalize 3’ CITE-seq libraries. GEMs were created on either a 10X Genomics Chromium Controller Genetic Analyzer (RRID:SCR_019326) or a Chromium X Single Cell Analyzer (RRID:SCR_024537). Either the Bio-Rad C-1000 (RRID:SCR_019688) or the Applied Biosystems SimpliAmp (RRID:SCR_023004) thermocyclers were used for library preparation.

### Single-Cell RNA QC and Sequencing

Quality and concentration of cDNA and final libraries were calculated from Agilent 2100 Bioanalyzer (RRID:SCR_018043) electrograms. Sequencing was performed on an Illumina NovaSeq 6000 Sequencing System (RRID:SCR_016387).

#### RNA Isolation

RNA was isolated from the cultured NK cells using the PureLink RNA Mini Kit (Invitrogen; Ref. 12183018A) following the manufacturer’s instructions. Cells were pelleted at 2000 g for 5 minutes. The media was aspirated, and cells were lysed in 600µL of the provided lysis buffer. Cells were manually homogenized using a 20 g needle (BD and Company; Model 305176) and a 3mL syringe (BD and Company; 309657) 10 times. The RNA was purified using the kit-provided columns and wash buffers. The RNA was eluted in 60µL of the provided RNase-free water.

### Quantitative PCR (qPCR)

For testing the relative amount of ERP gene mRNAs in isolated NK cells, qPCR assays were created with FAM-labeled probes for *NR4A2*, *FOS*, and *JUNB* (IDT). Each qPCR reaction also contained a 20x*YWHAZ*-VIC assay (Thermo Fisher Scientific; Gene ID: Hs03044281_g1; Cat. # 4331182) as a housekeeping gene control. The ERP gene assays used (*gene name*, *NCBI Reference Sequence number, amplicon location, Forward Primer, Reverse Primer, and FAM probe)*: *NR4A2, NM_006186.4,* 365-455, AGC CAT GCC TTG TGT TCA, TCT CCC GAA GAG TGG TAA CT, and FAM-TT GAG GCG A-ZEN-G GAC CCA TAC TGC-Iowa Black FQ; *FOS*, NM_005252.2, 56-163, CAG CGA ACG AGC AGT GA, ACA TCA TCG TGG CGG TTA G, and FAM-TC CTA CCC A-ZEN-G CTC TGC TCC ACA-Iowa Black FQ; and *JUNB*, *NM_002229.3, 281-376,* GCC CGG ATG TGC ACT AAA, TGT AGA GAG AGG CCA CCA G, FAM-AG CCC TTC T-ZEN-A CCA CGA CGA CTC ATA-Iowa Black FQ. The qPCR was performed with AgPath-ID One-step RT-PCR kit (Applied Biosystems; 4387391) following manufacturer recommendations for 20uL qPCR reaction volumes. YWHAZ-VIC was used at 1x, 1µL per reaction. Oligos were used at 500µM for primers and 250µM for probes, or 1µL from pre-mixed assays. Each well contained 2µL of isolated RNA and 5µL of UltraPure Distilled Water. Each time point was performed separately on MicroAmp 96-well plates (Applied Biosystems; 4346906) and covered with MicroAmp Optical Adhesive Film (Applied Biosystems; 4311971). A master mix of 2x Buffer, enzyme mix, and ddH2O plus 15% extra was made for all conditions (control RNA, TGF-β1 RNA, and no RNA) and primer mix (*NR4A2*-FAM/*YWHAZ*-VIC, *FOS*-FAM/*YWHAZ*-VIC, and *JUNB*-FAM/*YWHAZ*-VIC). The master mixes plus 10% were aliquoted for the three (3) conditions, and RNA or water, plus 10% was added. The master mix and RNA were aliquoted for each primer mix plus 5%. Finally, 20µL of each reaction mix was added to the wells of the qPCR plate. QuantStudio 7 Pro Real-Time PCR System, 96-well (Applied Biosystems) was used for the qPCR using the program: 50°C for 2 minutes, 95°C for 10 minutes, and 40 cycles of: 95°C for 15 seconds and 60°C for 1 minute.

## QUANTIFICATION AND STATISTICAL ANALYSIS

### Approach for Spectral Flow Analysis

Spectral flow cytometry results were analyzed using a combination of FlowJo v10.10.0 Software for Mac (RRID:SCR_008520), FlowSOM v3.0.18 (RRID:SCR_016899), Adobe Photoshop for Mac (RRID:SCR_014199), Microsoft Excel for Mac (RRID:SCR_016137).

Before concatenating the LiveCD45+CD14-CD3-CD56+ cell data from eight (8) healthy control endometrial biopsies, data was exported from FlowJo into Excel to generate the charts found in Figures S1C and S2B. This was to show that the eight (8) healthy controls had roughly the same percentage of CD56+ cells within each group of cells discussed in this paper. After concatenation, data was exported from FlowJo into Excel once again to illustrate the percentage of cell types found within the flow data set, as shown in Figure S1B.

FlowJo’s native t-distributed stochastic neighbor embedding (t-SNE) algorithm was used on the concatenated data to generate the initial t-SNE plots using the following parameters: BV421::CD103, BV711::CD49a, FITC::KIRS, PE/Cyanine7::CD39, and PE/Dazzles594::CD16. The 45 populations shown in Figure 1C were found by FlowSOM v3.0.18 using the previously generated t-SNE and the same fluor::marker parameters as inputs.

All phenotype, population, and heatmap t-SNE plots found in Figure 1 were produced using a combination of FlowJo’s Layout Editor and Cluster Explorer algorithm. The overlay plots, however, were created using Adobe Photoshop for Mac. A population t-SNE plot from Cluster Explorer was inserted into a Photoshop layer, while a phenotype t-SNE plot also from Cluster Explorer was inserted into the second layer. After ensuring the plots lined up perfectly, the Exclusion blending mode was applied to both layers. Both layers were then merged, and an Invert adjustment layer was applied to export the resulting plot.

Spectral flow cytometry analysis for the TGF-β incubation was performed using FlowJo software (v10.10.0). Single-cell suspensions were first gated on forward and side scatter properties to exclude debris. Doublets were excluded using SSC-H vs SSC-A gating. Viable cells were selected using LiveDead Aqua dye. NK cells were identified as CD3⁻CD14⁻CD19⁻CD56⁺ cells, and integrin expression (CD49a and CD103) was assessed within this population. Fluorescence minus one (FMO) controls were used to establish gating boundaries for positivity. Data are presented as the percentage of CD103⁺ cells within the NK cell population and median fluorescence intensity (MFI) of CD103 expression.

### Cell and Cluster Assignment by Thresholding

*AddModuleScore* was used with decidual NK signatures (below) corresponding to dNK1, dNK2, and dNK3 to calculate module scores. Generally, when using gene sets established in prior studies on similar but not identical tissue (namely decidua versus endometrium), it is unknown *a priori* which, if any, clusters correspond to previously published cell types, particularly cell types recovered from different tissues (decidua versus endometrium). It is also not known whether data under consideration are over- or under-clustered relative to procedures used to generate previously published modules; therefore it is also unknown whether more than one cluster corresponds to previously published cell types. We thus developed a procedure to assign one or more endometrial NK cell clusters to each of the three previously published gene sets representing decidual NK cells.

We first assumed that our data contained at least one cluster representing each of the three decidual cell types. We further assumed that two or more signatures would not describe a single cluster equally well, and therefore that any given cluster could represent at most a single signature. With these assumptions in mind, for each of the three signatures we ranked clusters in descending order of the mean module score, producing three lists in which each cluster was listed once. On the basis of the first assumption, the first cluster in each list was assigned to that signature. Remaining clusters were provisionally assigned to each signature according to their relative order in the three lists. For example, the second entry in one list was assigned to that signature and was thus excluded from assignment from any other cluster. Clusters that were listed in the same position in two lists were subject to a tie-breaking test described below. The result is a provisional list of clusters that could correspond to previously published NK cell types.

After provisional signature-cluster assignments, we then performed an iterative thresholding procedure to determine the maximum possible predictive power of signature scores in assigning individual cells to the clusters associated with each signature. For each signature, beginning with the first assigned cluster, we determined a threshold score value that minimized the absolute difference between the number of cells assigned to the cluster and below the threshold and the number of cells not assigned to the cluster and above the threshold. This is equivalent to jointly minimizing type I and type II errors: cells are regarded as false positives if they are above the threshold but not in the cluster, and false negatives if they are below the threshold but in the cluster. The minimization yielded a score value used as a threshold to then calculate the positive likelihood ratio, the negative likelihood ratio, and the odds ratio. These values represent measurements of the best possible performance of the signature for predicting inclusion in and exclusion from the cluster under consideration.

We then asked whether the predictive power of the scores increased or decreased by adding additional provisional clusters one at a time, moving down the ranked cluster list. For each additional cluster, we determined the revised threshold and recalculated the performance metrics. If inclusion of additional clusters resulted in an increase in performance (i.e. an improved odds ratio, regardless of whether it resulted from either an increase in the positive likelihood ratio and/or a decrease in the negative likelihood ratio), then additional clusters were counted as definitively assigned to the signature in question.

For clusters in the same position in two or more score lists, we compared the change in performance from including that cluster in one or the other signature. Clusters were definitively assigned to the signature for which they increased the odds ratio. If the odds ratios increased in both cases, the cluster was assigned to the signature for which its inclusion yielded the largest increase. If inclusion decreased the odds ratio of both scores, the cluster was not assigned any signature.

Note that whereas each signature was assigned at least one cluster on account of the first assumption above, not all clusters were assigned to signatures. This occurred for clusters whose addition to any signature decreased the odds ratio.

### Derivation of Gene Signatures

Decidual natural killer (dNK) cell gene modules were derived from differentially expressed genes defining dNK subsets (Vento-Tormo et al., Supplementary Table 7). DEGs were filtered by average log fold-change ≥0.5 and adjusted p-value ≤0.05, then ranked by descending average log fold-change. The top 15 genes for each subset were selected to generate dNK1, dNK2, and dNK3 module scores. Conventional natural killer (cNK) cell gene modules were derived from differentially expressed genes defining blood NK subsets (Yang et al., Supplementary Table 2). Peripheral blood natural killer (pbNK) cell gene modules derived from literature. TGF-β target gene modules were derived from curated lists of experimentally validated TGF-β transcriptional targets compiled from previously published studies.

### qPCR Analysis

Design and Analysis v2.8.0 software (RRID:SCR_026962) was used to analyze the qPCR results for Cq values. The ΔCq values used in Figure 6B were calculated by subtracting the Cq values of the target gene of each well by the YHMZ-VIC Cq values for the corresponding well. Two-tailed Student’s t-test was performed in MATLAB (RRID:SCR_001622).

### scRNA-Seq data processing and analysis

Reads from single-cell RNA-seq experiments of each healthy control (HC) library were pre-processed and aligned against the GRCh38 human reference and quantified using the Cell Ranger Single-Cell Software Suite v6.1.1 (RRID:SCR_017344) from 10x Genomics. Downstream analysis of the generated count matrix was completed using R v4.2.1 (RRID:SCR_001905) and Seurat v4.4.0 (RRID:SCR_007322), utilizing default parameters unless otherwise specified. Prior to subsequent analyses, background noise from ambient RNA was removed using SoupX v1.6.2 (RRID:SCR_019193). The overall contamination fraction (rho) was parameterized using the *autoEstCont* function to remove > 2% background contamination in each dataset. The SoupX-corrected count matrixes were then loaded into R using the *Read10X* function from which Seurat objects were created per dataset via the *CreateSeuratObject* function. From each dataset, cells with less than 200 features, and features with less than three (3) cells expressing them were excluded. Additionally, respective count matrices were filtered based on quality control (QC) metrics and thresholds (Supp table Sx and Supp fig. Sx). Doublets arising from the 10X sample loading procedure were detected and removed using Scrublet v0.2.3 (RRID:SCR_018098). The doublet rate per library was approximated using estimated rates provided by 10x Genomics.

Filtered count matrices were then merged into one Seurat object, normalized, and variance stabilized using regularized negative binomial regression via the *SCTransform* function (RRID:SCR_022146) provided by Seurat. We also regressed out the variation introduced by mitochondrial expression to prevent any confounding signal. Next, identification of principal components was performed using *RunPCA()*. Leveraging identified PCs, and while considering each individual library as a batch, the normalized dataset was corrected for batch effects and integrated using *Harmony* v1.2.0 (RRID:SCR_022206). The resulting top 20 Harmony-corrected dimensions were further used to construct a k-nearest neighbor (KNN) graph using *FindNeighbors()*. Cell clustering was subsequently performed using *FindClusters(),* which employs a shared nearest neighbor (SNN) modularity optimization-based clustering algorithm. The resulting cluster information was used as input into the uniform manifold approximation and projection algorithm, RunUMAP(), which further aided the visualization of cell manifolds in a low-dimensional space. We ran Seurat’s implementation of the Wilcoxon rank-sum test *FindMarkers()* to identify differentially expressed genes in each cluster. The expression of cluster-specific canonical markers was used to annotate each cell cluster.

### CITE-Seq data processing and analysis

Datasets from three healthy controls (HC10, HC12, and HC20) were further used for cell surface protein and transcriptomic data analyses. Pre-processing of these data was conducted as described previously. The RNA gene expression data were normalized using SCTransform() (as earlier described); however, the ADT counts were centered log-ratio (CLR) normalized. Normalized ADT counts were then scaled and centered using the *ScaleData* function. Principal Component Analysis (PCA) was conducted on both assays (RNA and ADT) using the *RunPCA* function with default parameters. We leveraged the *ElbowPlot()* and *DimHeatmaps()* to determine the number of principal components (PCs) to consider for further downstream analyses. Towards cell clustering, we found the k-nearest neighbors of each cell using the Seurat *FindNeighbors* function, which then constructed a shared nearest neighbor graph using the top 35 and top 20 dimensions for RNA and ADT assays, respectively. As the data constitute multiple modalities, we leveraged Seurat to define cellular states based on these two assays. We applied the weighted-nearest neighbor (WNN) analysis, an unsupervised framework to integrate the multiple data types measured within each cell, and to obtain a unified definition of cellular state based on the multiple modalities. For each cell, a set of modality weights was learned, which reflects the relative information content for each data type in that cell. This enabled the generation of a WNN graph that denotes the most similar cells in the dataset based on a weighted combination of protein and RNA similarities. To achieve this, we used the Seurat *FindMultiModalNeighbors* function, generating cell neighborhood graphs for further downstream analyses. We then performed cell clustering on the generated graphs and identified clusters using the Louvain algorithm (*FindCluster* function) at the default resolution. We utilized cluster information as input into the uniform manifold approximation and projection (UMAP) activated by the *RunUMAP* function which further aided the visualization of cell manifolds in a two-dimensional space, and as a weighted combination of RNA and protein data.

To identify differentially expressed (DE) genes between groups of cells, we used the *FindMarkers* function using MAST (RRID:SCR_016340). Results from differential gene expression analysis were used to annotate clusters based on the expression of top marker genes in the resulting clusters, and by leveraging feature plots (*FeaturePlot* function) of canonical cell type specific marker expression.

### Trajectory inference of single cells

Analysis of NK cell trajectories was performed using Monocle3 v1.3.4 (RRID:SCR_018685) on the HC18 Seurat object. Preprocessing was conducted as described previously. The HC18 dataset was converted to a Monocle3 dataset using *as.cell_data_set*(). Clustering information from prior analyses was maintained and leveraged for trajectory inference. Cells were then organized into trajectories using *SimplePPT* (simple principal tree algorithm), a reversed graph embedding (RGE) method implemented through the *learn_graph* function from Monocle3, at default parameters, and further visualized using the *plot_cells* function. For pseudotemporal ordering of cells according to their transcriptome similarity, we defined the root of the trajectory both programmatically by running *get_earliest_principal_node(),* and based on signatures in literature that define early residency, setting CD56^bright^ pbNK cells as the root of the trajectory and using the *order_cells* function to order the cells based on the set root. To identify differentially expressed genes (DEGs) that change as a function of pseudotime along the trajectory, we conducted a *graph_test()* using the Moran’s I test to detect significant genes showing correlation along the principal graph embedded in the low-dimensional space. We then grouped identified DEGs (q_value < 0.05) into modules to further elucidate cell fate specific genes and genes that are activated at different positions on the trajectory or cell cluster. Trajectory-variable genes were collected and clustered into modules that are co-expressed across cells using the *find_gene_modules* function. Furthermore, *aggregate_gene_expression()* was used to compute aggregate module scores within each group of cell types. The resulting aggregate module scores per cluster were plotted and visualized using the *pheatmap* function (RRID:SCR_016418). To provide further insights into the gene module dynamics along the trajectory, computed gene modules were passed to the *plot_cells* function and visualized as a superimposition on the UMAP, and along the trajectory. We also leveraged the *plot_genes_in_pseudotime* function to show specific gene dynamics as a function of pseudotime and across cell clusters. Additionally, the top 10 differentially expressed genes with q_value < 0.05 & morans_I > 0.1 per module were used to cluster genes based on their pseudotemporal expression patterns.

### Gene set enrichment analysis

Gene set enrichment analysis was performed using the FGSEA package (RRID:SCR_020938). Inputs included gene sets of interest and ranked differentially expressed gene lists created with MAST and the Seurat *FindMarkers* function. Resulting differentially expressed genes were ranked based on the log-fold change (or a combinatory metric of absolute log-fold change * log transformed p-values) and the ranked gene list used as input for the FGSEA R package (fast preranked GSEA), with a minimum set size of 15 genes and a maximum of 500 genes. We used the *plotEnrichment* function to visualize the resulting gene set enrichment. Analyzed gene sets were manually curated through literature review as well as analysis of publicly available data.

### Module scores

Module scores were calculated using the *AddModuleScore* function from Seurat. Respective feature plots were generated using *FeaturePlot()* with the features set as the calculated module scores from each gene program. Hypergeometric tests for gene overlaps between modules was conducted using the *GeneOverlap* R package (RRID:SCR_018419) at default parameters and spec/genome name hg19.gene.

