## Supplementary material for "A developmental program of early residency promotes the differentiation of divergent uterine NK cell subsets in humans": Supplemantary_Figures

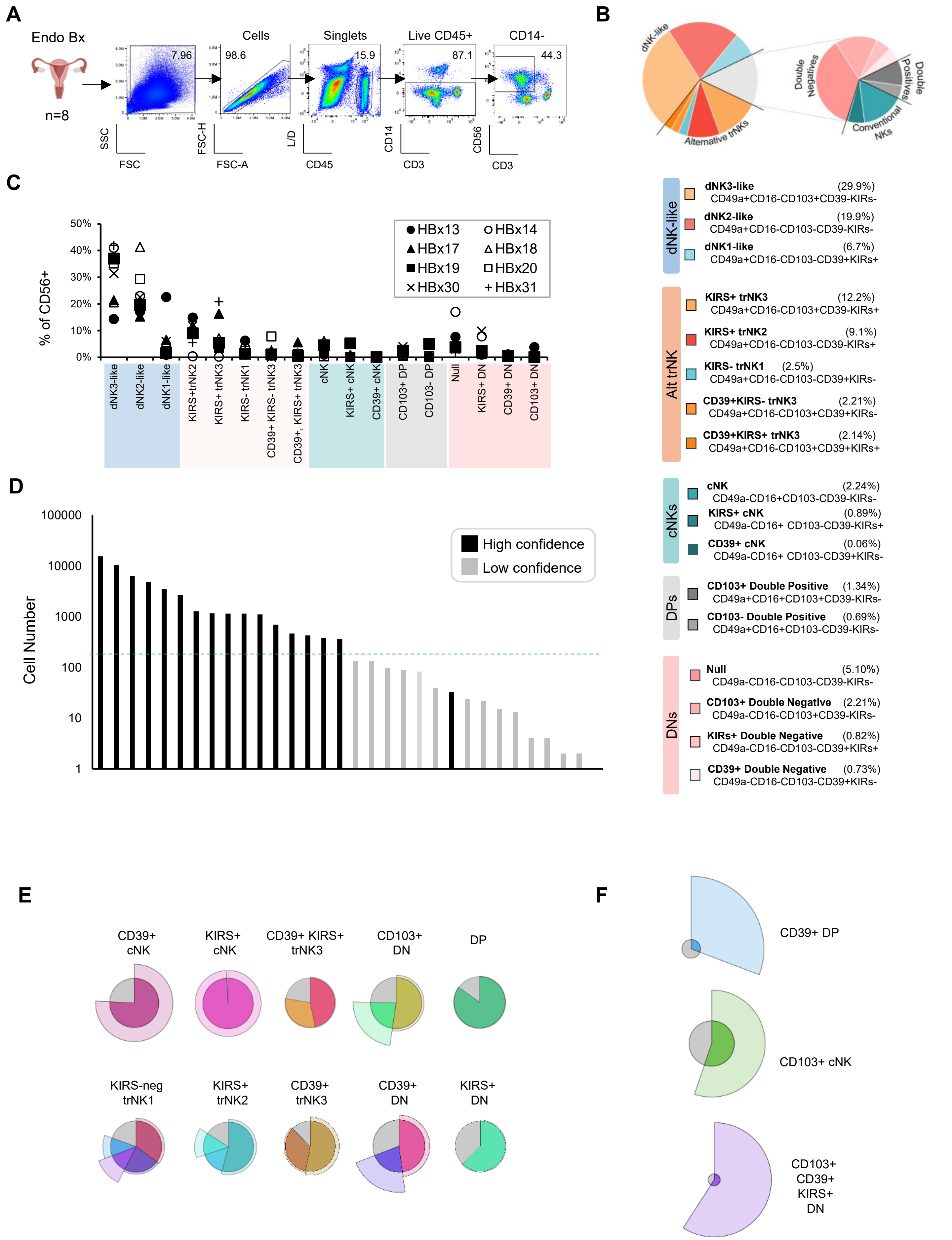
A
Endo Bx
n=8
Cells
Singlets
Live CD45+
CD14-
7.96
98.6
15.9
87.1
44.3
SSC
FSC
FSC-H
FSC-A
L/D
CD45
CD14
CD3
CD56
CD3
B
dNK-like
Alternative trNKs
Double Negatives
Double Positives
Conventional NKs
C
% of CD56+
HBx13
HBx14
HBx17
HBx18
HBx19
HBx20
HBx30
HBx31
dNK3-like
dNK2-like
dNK1-like
KIRS+trNK2
KIRS+ trNK3
KIRS- trNK1
CD39+ KIRS- trNK3
CD39+, KIRS+ trNK3
cNK
KIRS+ cNK
CD39+ cNK
CD103+ DP
CD103- DP
Null
KIRS+ DN
CD39+ DN
CD103+ DN
dNK-like
dNK3-like (29.9%)
CD49a+CD16-CD103+CD39-KIRs-
dNK2-like (19.9%)
CD49a+CD16-CD103-CD39-KIRs-
dNK1-like (6.7%)
CD49a+CD16-CD103-CD39+KIRs+
Alt trNK
KIRS+ trNK3 (12.2%)
CD49a+CD16-CD103+CD39-KIRs+
KIRS+ trNK2 (9.1%)
CD49a+CD16-CD103-CD39-KIRs+
KIRS- trNK1 (2.5%)
CD49a+CD16-CD103-CD39+KIRs-
CD39+KIRS- trNK3 (2.21%)
CD49a+CD16-CD103+CD39+KIRs-
CD39+KIRS+ trNK3 (2.14%)
CD49a+CD16-CD103+CD39+KIRs+
cNKs
cNK (2.24%)
CD49a-CD16+CD103-CD39-KIRs-
KIRS+ cNK (0.89%)
CD49a-CD16+ CD103-CD39-KIRs+
CD39+ cNK (0.06%)
CD49a-CD16+ CD103-CD39+KIRs-
DPs
CD103+ Double Positive (1.34%)
CD49a+CD16+CD103+CD39-KIRs-
CD103- Double Positive (0.69%)
CD49a+CD16+CD103-CD39-KIRs-
DNs
Null (5.10%)
CD49a-CD16-CD103-CD39-KIRs-
CD103+ Double Negative (2.21%)
CD49a-CD16-CD103+CD39-KIRs-
KIRs+ Double Negative (0.82%)
CD49a-CD16-CD103-CD39+KIRs+
CD39+ Double Negative (0.73%)
CD49a-CD16-CD103-CD39+KIRs-
D
Cell Number
High confidence
Low confidence
E
CD39+ cNK
KIRS+ cNK
CD39+ KIRS+ trNK3
CD103+ DN
DP
KIRS-neg trNK1
KIRS+ trNK2
CD39+ trNK3
CD39+ DN
KIRS+ DN
F
CD39+ DP
CD103+ cNK
CD103+ CD39+ KIRS+ DN

Fig. S1

**Figure S1. Flow cytometric assessment of high and low confidence phenotypes among human uterine NK cells.**

(A-F) Eight human endometrial biopsies were analyzed with flow cytometry.

(A) Experimental schematic and gating strategy for analysis of human endometrial NK cells. Sequential gates are shown as indicated on a single representative sample of human endometrium after enzymatic digestion. Name above plot indicates parent gate. Live, CD45+ CD14- CD3- CD56+ singlets from each sample (n=8) were then concatenated in FlowJo for analysis in Figure 1.

(B) Frequency of 17 high confidence phenotypes (n=51,395 cells). Out of 32 potential phenotypes, 31 phenotypes were detected by manual gating and determined as high confidence after FlowSOM analysis.

(C) Frequency of high confidence NK cell phenotypes for individual samples (n=8).

(D) Number of cells in each of 31 detected phenotypes by manual gating, in descending order. High and low confidence phenotypes are indicated. Hashed line indicates cell number threshold (~100 cells) above which there is concordance of manual gating and FlowSOM results.

(E) Polar area charts of 10 of 17 high confidence phenotypes to complement Figure 1D.

(F) Representative polar area charts of low confidence phenotypes. Note that most cells in a given population are included in the shaded area outside of the central pie (phenotype), indicating that the majority of cells in that population do not belong to the given phenotype. Central pie is not drawn to scale with prior panel to accommodate size of shaded area outside of central pie.

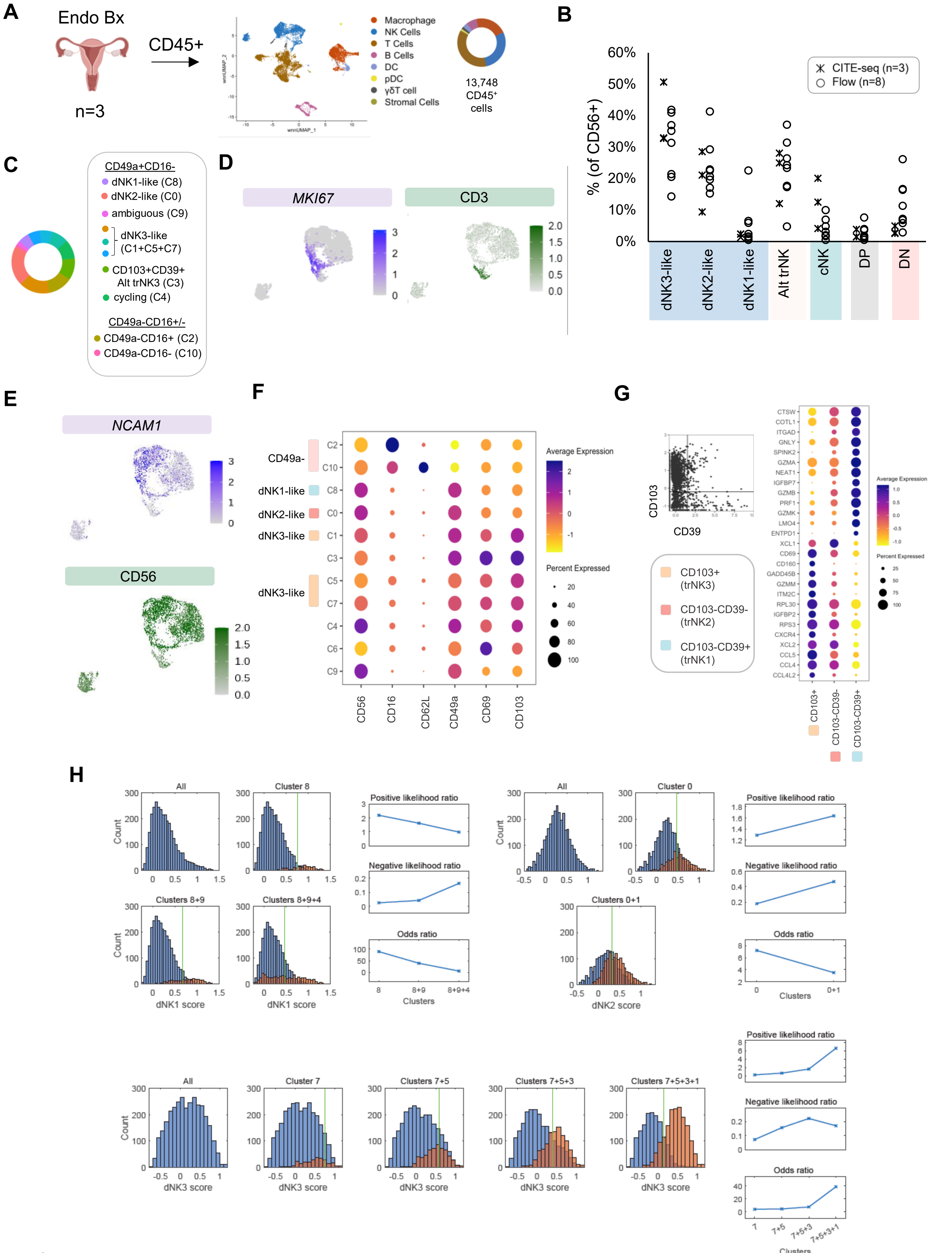

A
Endo Bx
n=3
CD45+
Macrophage
NK Cells
T Cells
B Cells
DC
pDC
γδT cell
Stromal Cells
13,748 CD45+ cells
B
CITE-seq (n=3)
Flow (n=8)
% (of CD56+)
dNK3-like
dNK2-like
dNK1-like
Alt trNK
cNK
DP
DN
C
CD49a+CD16-
dNK1-like (C8)
dNK2-like (C0)
ambiguous (C9)
dNK3-like (C1+C5+C7)
CD103+CD39+ Alt trNK3 (C3)
cycling (C4)
CD49a-CD16+/-
CD49a-CD16+ (C2)
CD49a-CD16- (C10)
D
MKI67
CD3
E
NCAM1
CD56
F
CD49a-
dNK1-like
dNK2-like
dNK3-like
Average Expression
Percent Expressed
CD56
CD16
CD62L
CD49a
CD69
CD103
G
CD103
CD39
CD103+ (trNK3)
CD103-CD39- (trNK2)
CD103-CD39+ (trNK1)
CD103+
CD103-CD39-
CD103-CD39+
H
All
Cluster 8
Clusters 8+9
Clusters 8+9+4
dNK1 score
Count
Positive likelihood ratio
Negative likelihood ratio
Odds ratio
Clusters
Cluster 0
Clusters 0+1
dNK2 score
Cluster 7
Clusters 7+5
Clusters 7+5+3
Clusters 7+5+3+1
dNK3 score

Fig. S2

**Figure S2. Surface phenotypes and transcriptional profile of CD56+ NK cells in three secretory phase endometrial biopsies**.

(A-H) CITE-seq analysis of 3,957 CD56+ NK cells selected from 13,748 CD45+ immune cells enriched from secretory phase biopsies (n=3). Data were aggregated for analysis.

(A) Experimental schematic of endometrial biopsies and sorting of CD45+ immune cells. NK cells were selected and re-clustered for analysis in Figure 2.

(B) No statistically significant difference in the frequency of NK subsets in samples analyzed by flow cytometry (n=8) (see Figure 1) vs. CITE-seq (n=3) (see Figure 2). Major groups of phenotypes are shown.

(C) Distribution of 11 clusters of 3,957 CD56+ endometrial NK cells.

(D) Feature plots of expression of *MKI67* (to identify cycling cells; cluster 4) (left) and CD3 (to identify NKT cells; cluster 6, excluded) (right).

(E) Feature plots of *NCAM1* and CD56 expression

(F) Bubble plot of surface protein expression.

(G) Expression of top differentially expressed genes between CD103+ (trNK3), CD103-CD39- (trNK2) and CD103-CD39+ (trNK1) cells.

(H) Cluster-signature pair assignments and optimum threshold determination. For each signature, clusters ranked by mean score are sequentially tested to determine the odds ratio after determining the optimum threshold for maximal prediction of the inclusion (and exclusion) of cells in (and out) of the cluster (see STAR Methods). The set of clusters that maximizes the odds ratio is assigned to the signature. For dNK1 and dNK2, a single cluster maximizes the odds ratio after thresholding, whereas additional clusters decrease the odds ratio and other performance metrics (positive and negative likelihood ratios). For dNK3, inclusion of four clusters maximizes performance.

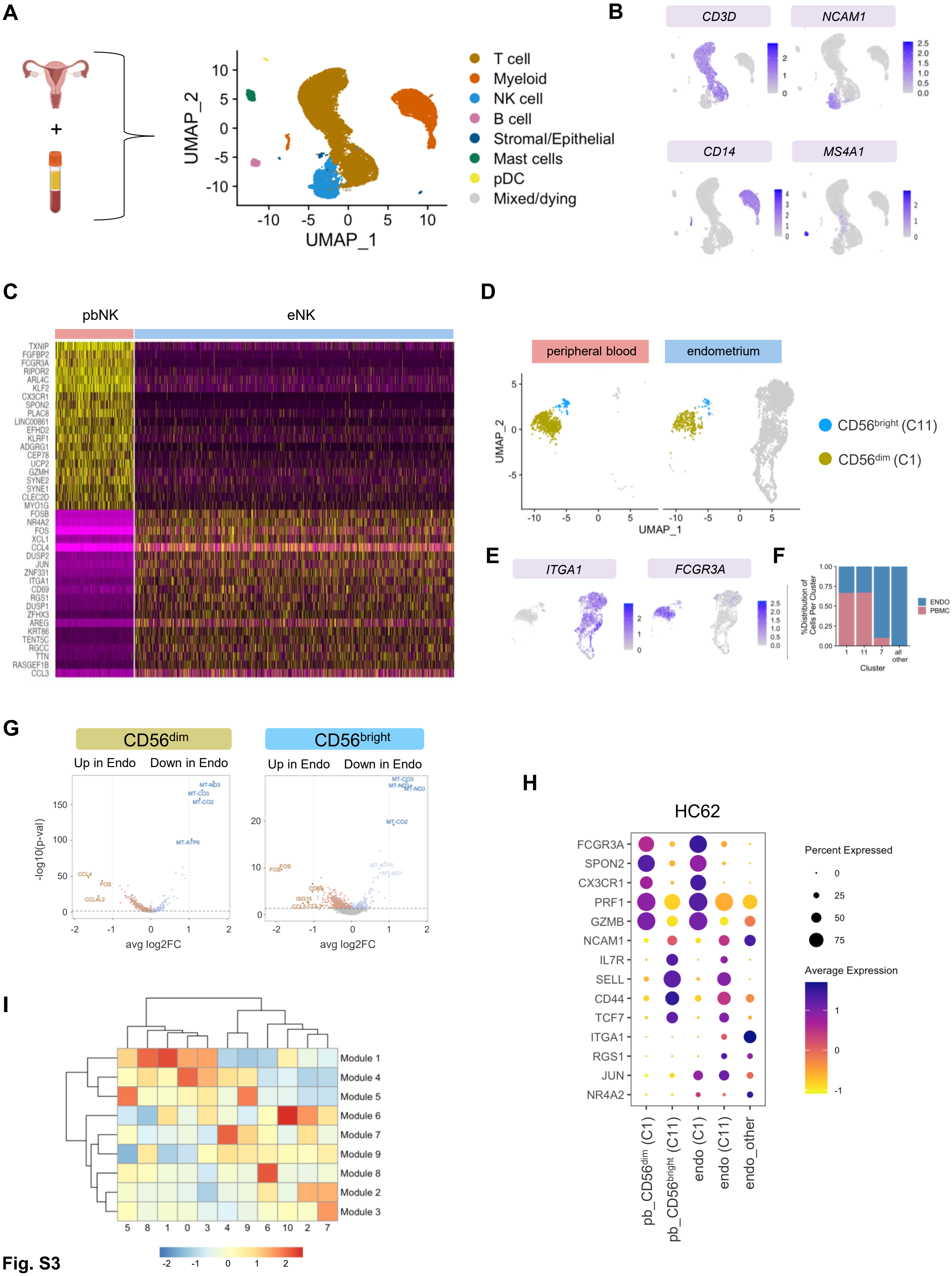
A
UMAP_2
UMAP_1
T cell
Myeloid
NK cell
B cell
Stromal/Epithelial
Mast cells
pDC
Mixed/dying
B
CD3D
NCAM1
CD14
MS4A1
C
pbNK
eNK
TXNIP
FGFBP2
FCGR3A
RIPOR2
ARL4C
KLF2
CX3CR1
SPON2
PLAC8
LINC00861
EFHD2
KLRF1
ADGRG1
CEP78
UCP2
GZMH
SYNE2
SYNE1
CLEC2D
MYO1G
FOSB
NR4A2
FOS
XCL1
CCL4
DUSP2
JUN
ZNF331
ITGA1
CD69
RGS1
DUSP1
ZFHX3
AREG
KRT86
TENT5C
RGCC
TTN
RASGEF1B
CCL3
D
peripheral blood
endometrium
UMAP_2
UMAP_1
CD56bright (C11)
CD56dim (C1)
E
ITGA1
FCGR3A
F
%Distribution of Cells Per Cluster
Cluster
ENDO
PBMC
all other
G
CD56dim
CD56bright
Up in Endo
Down in Endo
-log10(p-val)
avg log2FC
MT-ND3
MT-CO3
MT-CO2
MT-ATP6
MT-ND1
CCL4
FOS
CCL4L2
CD69
ISG15
CCL5
CCL3
H
HC62
FCGR3A
SPON2
CX3CR1
PRF1
GZMB
NCAM1
IL7R
SELL
CD44
TCF7
ITGA1
RGS1
JUN
NR4A2
Percent Expressed
Average Expression
pb_CD56dim (C1)
pb_CD56bright (C11)
endo (C1)
endo (C11)
endo_other
I
Module 1
Module 4
Module 5
Module 6
Module 7
Module 9
Module 8
Module 2
Module 3

Fig. S3

**Figure S3. Matched blood and endometrium schematic and validation approach**

(A-C,I) scRNA-seq of 2,495 natural killer cells re-clustered from endometrial and peripheral blood CD45+ immune cells. Samples were matched from one individual (HC18) and collected within 24h during the secretory phase of the menstrual cycle. eNK = endometrial NK; pbNK = peripheral blood NK.

(A-B) Schematic of experimental design (A) and identification of NK cells in aggregated CD45+ dataset (B). NK cells were selected and re-clustered for analysis in Figures 3&4.

(C) Comparison of peripheral blood NK cells with endometrial NK cells. Top differentially expressed genes are shown.

(D-H) scRNA-seq of 5,119 natural killer cells re-clustered from endometrial and peripheral blood CD45+ immune cells. Samples were matched from one individual (HC62) and collected within 24h during the secretory phase of the menstrual cycle. eNK = endometrial NK; pbNK = peripheral blood NK.

(D) UMAP embeddings of endometrial and peripheral blood NK cells, split by sample.

(E) Reciprocal expression of *FCGR3A* and *ITGA1* demarcates circulating peripheral blood NK cells from tissue-resident endometrial NK cells.

(F) Frequency of pbNK and/or eNK cells in each cluster.

(G) Comparison of peripheral blood and endometrial NK cells for indicated clusters.

(H) Expression of marker genes of peripheral blood $CD56^{dim}$ and $CD56^{bright}$ cells, along with genes associated with tissue residency in pbNK and endoNK.

(I) Heatmap of trajectory module scores, ordered by hierarchical clustering.

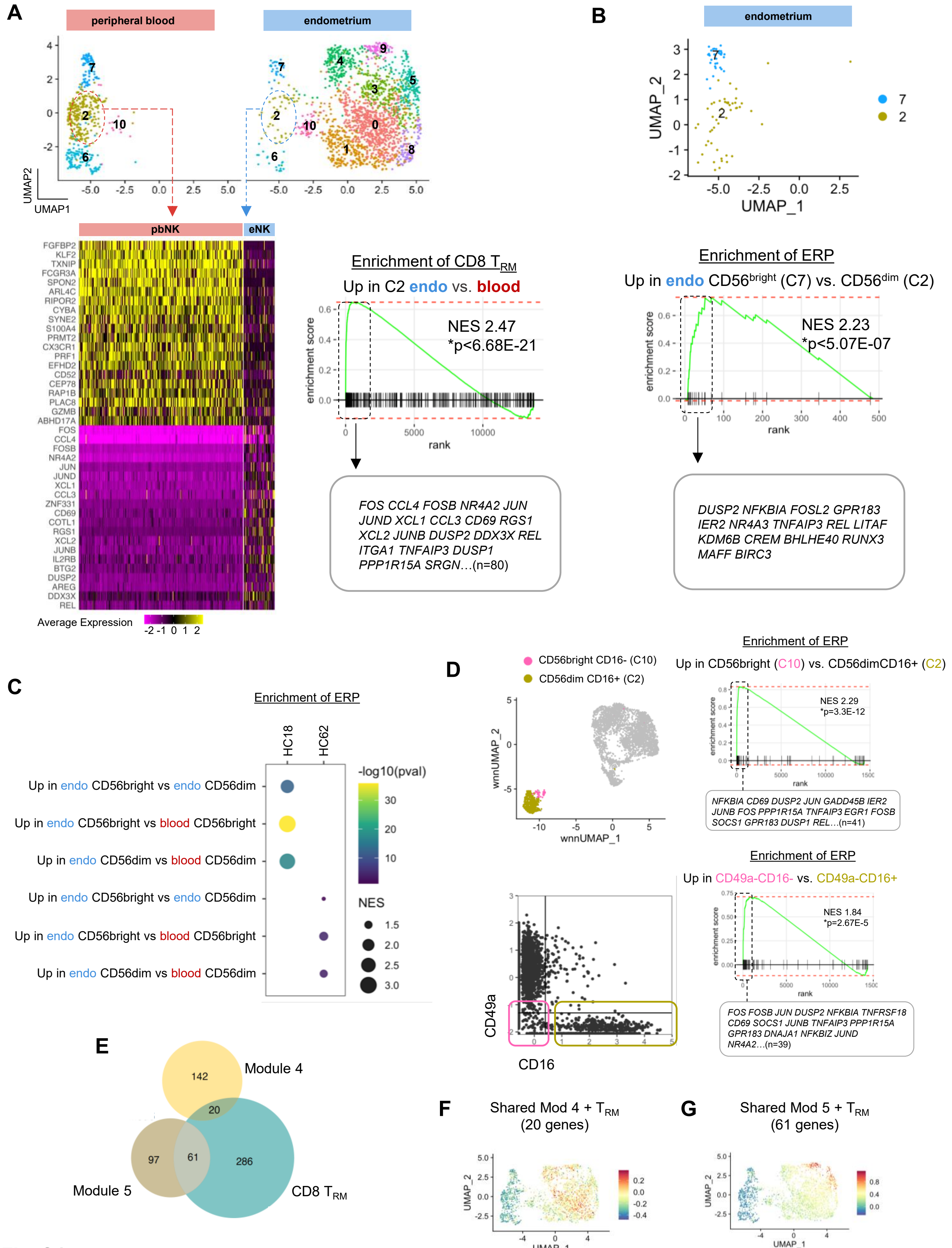
A
peripheral blood
endometrium
UMAP2
UMAP1
pbNK
eNK
FGFBP2
KLF2
TXNIP
FCGR3A
SPON2
ARL4C
RIPOR2
CYBA
SYNE2
S100A4
PRMT2
CX3CR1
PRF1
EFHD2
CD52
CEP78
RAP1B
PLAC8
GZMB
ABHD17A
FOS
CCL4
FOSB
NR4A2
JUN
JUND
XCL1
CCL3
ZNF331
CD69
COTL1
RGS1
XCL2
JUNB
IL2RB
BTG2
DUSP2
AREG
DDX3X
REL
Average Expression
Enrichment of CD8 T_RM
Up in C2 endo vs. blood
NES 2.47
*p<6.68E-21
enrichment score
rank
FOS CCL4 FOSB NR4A2 JUN JUND XCL1 CCL3 CD69 RGS1 XCL2 JUNB DUSP2 DDX3X REL ITGA1 TNFAIP3 DUSP1 PPP1R15A SRGN…(n=80)
B
endometrium
UMAP_2
UMAP_1
Enrichment of ERP
Up in endo CD56bright (C7) vs. CD56dim (C2)
NES 2.23
*p<5.07E-07
enrichment score
rank
DUSP2 NFKBIA FOSL2 GPR183 IER2 NR4A3 TNFAIP3 REL LITAF KDM6B CREM BHLHE40 RUNX3 MAFF BIRC3
C
Enrichment of ERP
HC18
HC62
Up in endo CD56bright vs endo CD56dim
Up in endo CD56bright vs blood CD56bright
Up in endo CD56dim vs blood CD56dim
Up in endo CD56bright vs endo CD56dim
Up in endo CD56bright vs blood CD56bright
Up in endo CD56dim vs blood CD56dim
-log10(pval)
NES
D
CD56bright CD16- (C10)
CD56dim CD16+ (C2)
wnnUMAP_2
wnnUMAP_1
Enrichment of ERP
Up in CD56bright (C10) vs. CD56dimCD16+ (C2)
NES 2.29
*p=3.3E-12
enrichment score
rank
NFKBIA CD69 DUSP2 JUN GADD45B IER2 JUNB FOS PPP1R15A TNFAIP3 EGR1 FOSB SOCS1 GPR183 DUSP1 REL…(n=41)
CD49a
CD16
Enrichment of ERP
Up in CD49a-CD16- vs. CD49a-CD16+
NES 1.84
*p=2.67E-5
enrichment score
rank
FOS FOSB JUN DUSP2 NFKBIA TNFRSF18 CD69 SOCS1 JUNB TNFAIP3 PPP1R15A GPR183 DNAJA1 NFKBIZ JUND NR4A2…(n=39)
E
Module 4
Module 5
CD8 T_RM
142
20
97
61
286
F
Shared Mod 4 + T_RM
(20 genes)
UMAP_2
UMAP_1
G
Shared Mod 5 + T_RM
(61 genes)
UMAP_2
UMAP_1

Fig. S4

**Figure S4. ERP enrichment in CD56bright endometrial founder NK cells across tissue samples.**

(A-B, E-G) scRNA-seq results of 2,495 natural killer cells re-clustered from endometrial and peripheral blood CD45+ immune cells. Samples were matched from one individual (HC18) and collected within 24h during the secretory phase of the menstrual cycle.

(A) Enrichment of the CD8 $T_{RM}$ gene signature in HC18 cluster 2 endometrial CD56$^{dim}$ cells vs. cluster 2 peripheral blood CD56$^{dim}$ cells. Heat map of top differentially expressed genes between pbNK and eNK cells within CD56$^{dim}$ cluster 2 (left). Gene set enrichment analysis of CD8 $T_{RM}$ signature in genes up in endometrial CD56$^{dim}$ cells vs. cluster 2 peripheral blood CD56$^{dim}$. NES = normalized enrichment score. Leading edge genes are shown in the box.

(B) Enrichment of the ERP in genes differentially upregulated in endometrial CD56$^{bright}$ (C7) NK cells versus endometrial CD56$^{dim}$ (C2) NK cells in HC18. NES = normalized enrichment score. Leading edge genes are shown in the box.

(C-D) Validation of the ERP in multiple samples. scRNA-seq results of natural killer cells re-clustered from matched endometrial and peripheral blood CD45+ immune cells in HC18 and in HC62 (C). CITE-seq results of endometrial natural killer cells aggregated from three individuals (HC10, HC12, HC20).

(C) ERP enrichment in endometrial vs. peripheral blood NK cells and in endometrial CD56$^{bright}$ vs. CD56$^{dim}$ cells in two individuals (HC18, HC62). NES = normalized enrichment score.

(D) ERP enrichment in endometrial CD56$^{bright}$ vs. CD56$^{dim}$ cells selected in two different ways in aggregated CITE-seq data from three individuals (HC10, HC12, HC20). Cluster 10 cells with a CD56$^{bright}$ CD49a- CD16- phenotype and transcriptomic profile were compared to Cluster 2 cells with a CD56$^{dim}$ CD49a- CD16+ phenotype (top left). ERP enrichment analysis in genes differentially upregulated in C10 (CD56$^{bright}$) vs. C2 (CD56$^{dim}$) (top right). Endometrial NK cells were selected from CITE-seq data based on expression of CD49a and CD16 as shown (bottom left). ERP enrichment analysis was performed in genes differentially upregulated in C10 (CD49a-CD16-) vs. C2 (CD49a-CD16-) (top right).

(E) Overlap of developmental trajectory modules 4 and 5 derived from trajectory analysis of peripheral blood and endometrial NK cells in HC18 (Figure 4) with the CD8 $T_{RM}$ reference gene set. Analysis indicates that 20 genes from trajectory module 4 and 61 genes from trajectory module 5 are shared with genes that increase early in CD8 $T_{RM}$ differentiation.

(F-G) Module score expression of genes shared between the CD8 $T_{RM}$ reference signature and trajectory modules 4 (F) and 5 (G).

**A**

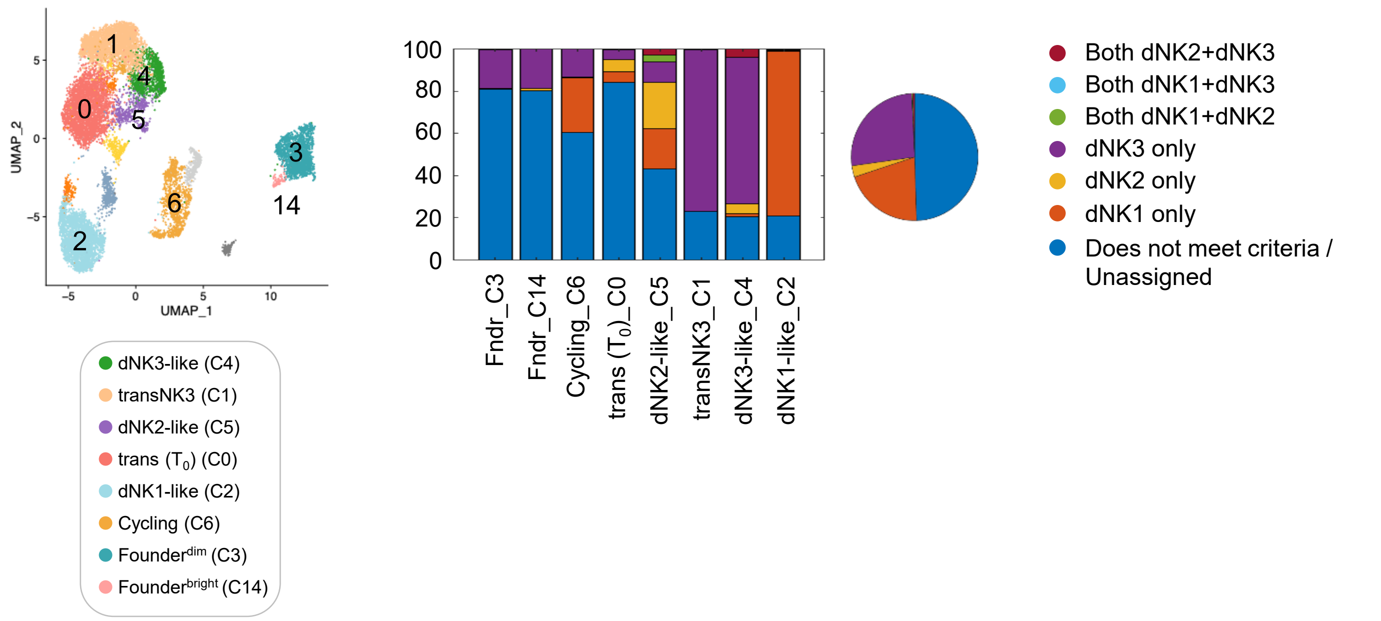

**B**

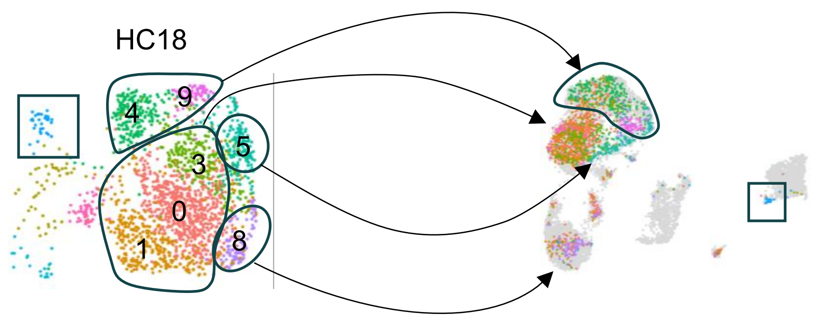

| | HC18 | 6HCAgg |
|---|---|---|
| dNK3-like | C4+C9 | C1+C4 |
| dNK2-like | C5 | C5 |
| dNK1-like | C8 | C2 |
| Transitional ($T_0$) | C0+C1+C3 | C0 |
| Founder (CD56[bright]) | C7 | C14 |

**C**

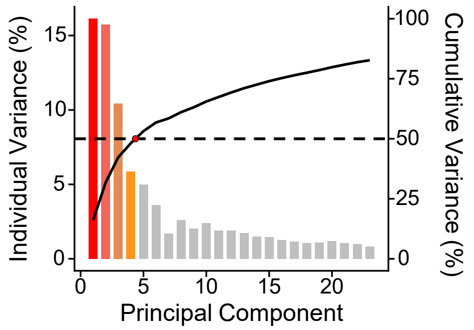

**D**

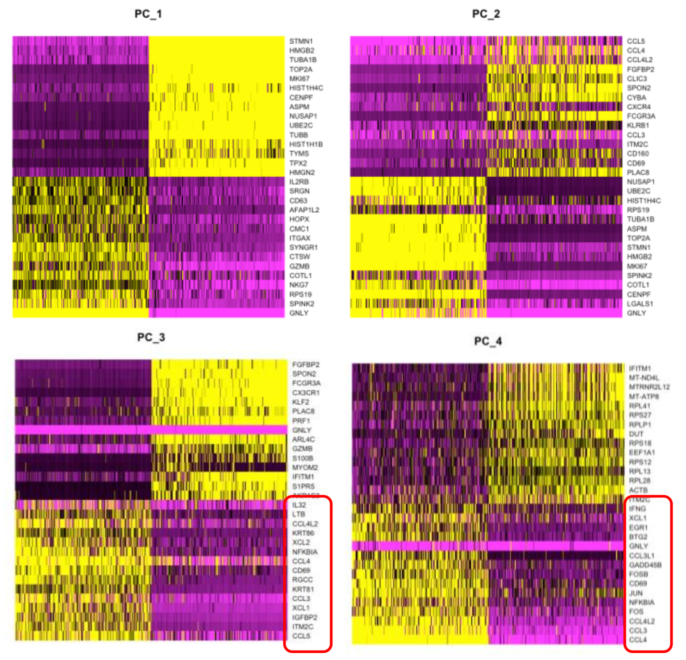

**E**

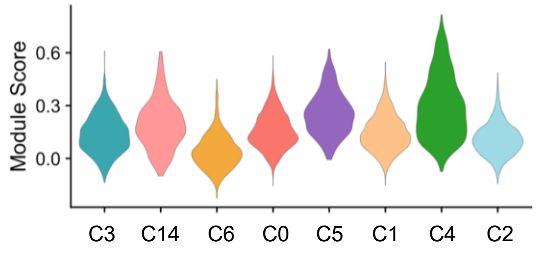

Fig. S5

**Figure S5. Contribution of the ERP to eNK transcriptional variability and differentiation of mature endometrial NK cells.**

(A-E) scRNA-seq of 14,349 natural killer cells re-clustered from 41,722 CD45+ immune cells enriched from secretory phase endometrium of six healthy control volunteers. Libraries were aggregated for analysis.

(A) Frequency of cell states within a cluster based on reference gene signature expression.

(B) Barcode mapping of HC18 identifies heterogeneity with transitional ($T_0$) cluster. Left UMAP indicates endometrial NK cell states after clustering with matched peripheral blood NK cells (Figure 3). Right UMAP portrays all endometrial NK cells aggregated from six healthy control volunteers. Cells from HC18 are colored by cell state assignment from aggregation with peripheral blood.

(C) Scree plot of variance for principal components in the dataset. Note that the top four principal components (red to orange) explain 50% of the variation in the data set (cumulative variance).

(D) Heatmap of eigenvalues for genes contributing to each principal component. Top four principal components are shown. Red box indicates subsets of ERP genes.

(E) ERP module score for each cluster.

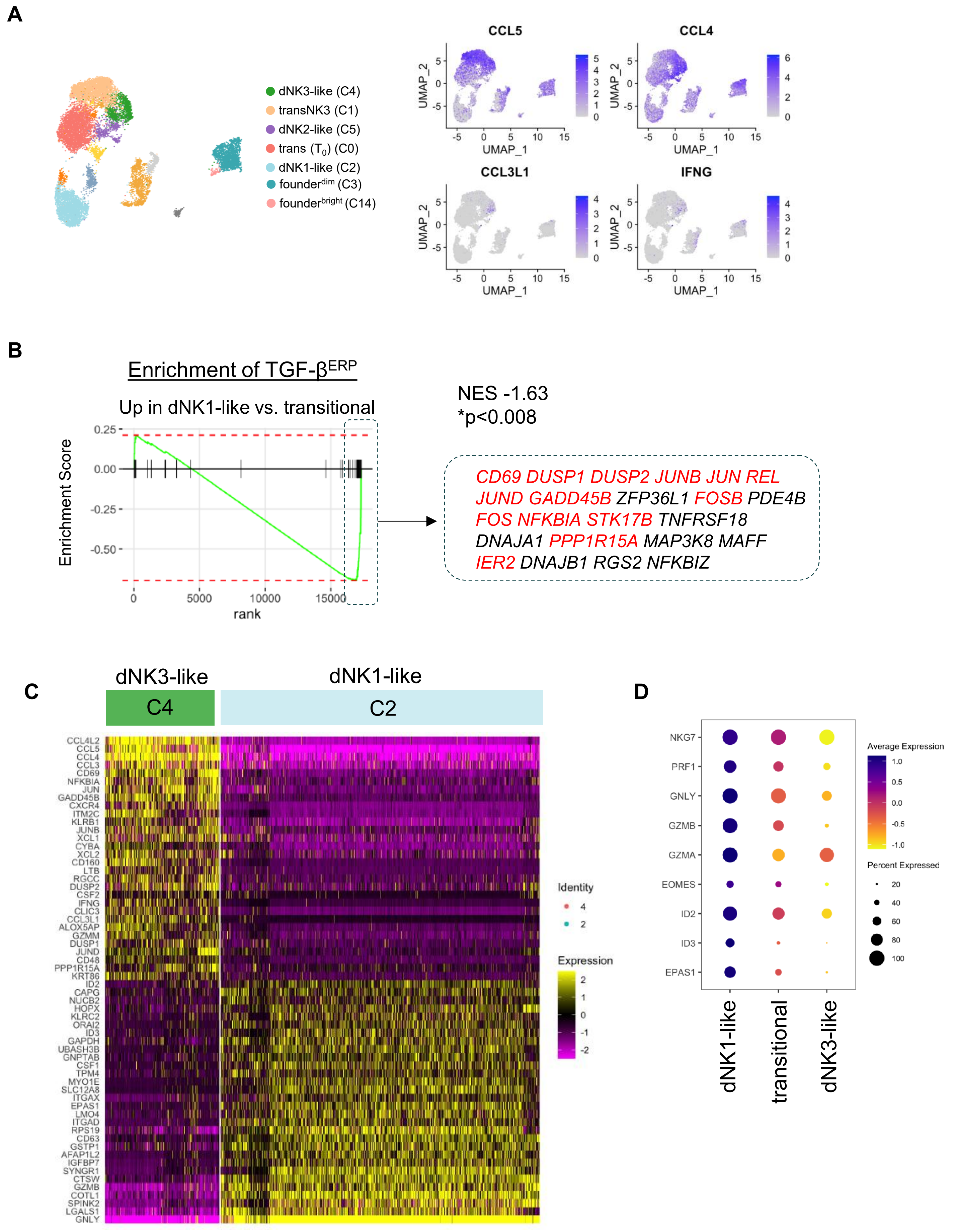
A
dNK3-like (C4)
transNK3 (C1)
dNK2-like (C5)
trans (T0) (C0)
dNK1-like (C2)
founderdim (C3)
founderbright (C14)
CCL5
CCL4
CCL3L1
IFNG
UMAP_1
UMAP_2
B
Enrichment of TGF-βERP
Up in dNK1-like vs. transitional
NES -1.63
*p<0.008
Enrichment Score
rank
CD69 DUSP1 DUSP2 JUNB JUN REL
JUND GADD45B ZFP36L1 FOSB PDE4B
FOS NFKBIA STK17B TNFRSF18
DNAJA1 PPP1R15A MAP3K8 MAFF
IER2 DNAJB1 RGS2 NFKBIZ
C
dNK3-like
dNK1-like
C4
C2
Identity
Expression
D
Average Expression
Percent Expressed
NKG7
PRF1
GNLY
GZMB
GZMA
EOMES
ID2
ID3
EPAS1
dNK1-like
transitional
dNK3-like

Fig. S6

**Figure S6. Differences in TGF-β responses between dNK1-like and dNK3-like eNKs.**

(A) Expression of chemokine ligand genes in endometrial NK cells. Note higher expression in dNK3-like and transitional NK3 cells with reduced or absent expression of selected effector genes in dNK1-like cells.

(B) Downregulation of TGF-$\beta^{ERP}$ in dNK1-like cells versus transitional cells. Top leading edge genes are shown in the box. Genes highlighted in red are part of the $ERP^{NK3}$ program, thus suggesting that the selective downregulation of these genes in dNK1-like cells may be due to antagonism of TGF-β signals.

(C) Differential expression analysis of dNK3-like and dNK1-like cells.

(D) Expression of genes known to be repressed by TGF-β in dNK1-like cells.
